# Structural and Thermodynamic Determinants of Anti-PGE_2_ mAb Specificity

**DOI:** 10.64898/2025.12.10.693102

**Authors:** Mitsuaki Sugahara, Hiromichi Saino, Michiyo Takehira, Yuko Kurahashi, Sae Aoyama-Sasabe, Katsuhide Yutani, Naoko Takahashi, Koji Takio, Hideo Ago, Masaki Yamamoto, Masashi Miyano, Shozo Yamamoto

## Abstract

We investigated the molecular determinants underlying the highly specific recognition of prostaglandin (PG) E_2_ and its structural analogues by an anti-PGE_2_ monoclonal antibody (mAb) and its Fab fragment using high-resolution X-ray crystallography coupled with detailed thermodynamic analyses, including conditions that mimic macromolecular crowding. Crystal structures of the Fab fragment in complex with PGE_2_, PGE_1_, and ONO5 were determined at 1.9 Å, 1.7 Å, and 1.7 Å resolution, respectively, and the antigen-free Fab structure was solved at 2.0 Å. The Fab accommodates the L-shaped PG scaffold within a deep antigen-binding crevasse formed between the β-sheet frameworks of the VH and VL domains, where ligand recognition is mediated by multiple water-bridged hydrogen bonds and extensive hydrophobic contacts, including putative CH–O and CH–π interactions. Comparison of antigen-free and ligand-bound structures revealed compensatory conformational rearrangements accompanied by reorganization of bound water networks.

Upon PG binding, a short 3_10_ helix forms at the tip of CDR-H1, stabilized by a cluster of hydrogen bonds and capping water molecules. This helix formation is associated with rotation of the VL domain and an increased Fab elbow angle. The CDR-H1 loop undergoes a flip-in transition in which Phe29H inserts into a hydrophobic cavity while Glu31H is extruded toward solvent, accompanied by π–π stacking between Tyr27H and Tyr32H. These changes collectively reduce hydrophobic surface area and promote enthalpy-driven ligand binding. The flexible α-chain of PGE_1_, lacking the C5–C6 double bond, adopts altered conformations that disrupt C1-carboxyl interactions with three aromatic residues and lead to closure of a secondary water channel, consistent with subtle but meaningful differences in binding thermodynamics and crystallographic temperature factors relative to PGE_2_ and ONO5.

Isothermal titration calorimetry (ITC) revealed high-affinity, enthalpy-driven PG binding to the Fab, in agreement with differential scanning calorimetry (DSC), which showed increased thermal stability upon complex formation. Under macromolecular crowding conditions induced by high bovine serum albumin (BSA) concentrations, PG binding to the intact mAb—but not to the Fab—exhibited improved discrimination between PGE_2_ and PGE_1_, suggesting that reduced entropy–enthalpy compensation under crowding amplifies subtle structural mismatches in ligand binding.

## Introduction

Prostaglandin (PG) E□ is a pivotal lipid mediator that exerts diverse physiological and pathophysiological effects, including roles in reproduction, neuronal function, maintenance of gastric mucosal integrity, smooth muscle tone, immune regulation, inflammation, pain, fever, and cancer progression (1–4). PGE□ is biosynthesized from arachidonic acid via PGH□ through cyclooxygenase activity, followed by conversion through multiple PGE□ synthases. It is subsequently inactivated by 15-hydroxyprostaglandin dehydrogenase and 15-oxoprostaglandin 13-reductase (5–8). The biological actions of PGE□ are mediated by its interaction with four G protein–coupled receptors, EP1–EP4 (9, 10).

Quantification of PGE□ is essential in biological research and clinical diagnostics, and many commercial ELISA kits are available capable of detecting concentrations from 10 pg to 10 μg. However, their specificity toward related prostanoids—including PGD□ and PGF□α—varies considerably among platforms. Some assays exhibit substantial cross-reactivity, whereas others show more than tenfold selectivity for PGE□ over PGF□α. Despite their widespread use, the structural basis for antigen recognition in these systems has not been elucidated. Previously, a mouse monoclonal antibody highly specific for PGE□ was developed (11). This antibody displays less than 10% cross-reactivity with PGE□ and PGF□α and less than 1% with other prostanoids such as PGD□, PGA□, PGB□, and thromboxane B□, but binds comparably to the PGE□ analogue 9-chloro-PGF□β (ONO5). Such high specificity is valuable for probing PGE□ function under pathological conditions (12), particularly given the distinct receptor-binding profiles and therapeutic applications of PGE□ and PGE□ (6, 10).

Understanding how antibodies discriminate subtle chemical differences among prostanoids is also relevant to ligand design in drug discovery.

To delineate the molecular mechanism underlying the remarkable specificity of antigen recognition by this anti-PGE□ monoclonal antibody, we conducted structural and thermodynamic analyses of its Fab fragment and intact mAb in complex with PGE□ and its analogues. Production of the Fab was optimized relative to previous protocols (13) to obtain sufficient quantities of highly purified material suitable for crystallography and calorimetric analyses. We determined high-resolution crystal structures of the Fab in the antigen-free form and in complex with PGE□, PGE□, and ONO5 (Fig. 1). Comparisons among these structures revealed several key features:

1. A large, deep antigen-binding crevasse between the VH and VL domains that fully accommodates the prostaglandin scaffold.
2. Ligand-induced conformational rearrangements, including a flip-flop transition in which hydrophobic and hydrophilic side chains exchange positions, formation of a short 3□□ helix stabilized by multiple water-mediated hydrogen bonds, and a small but significant rotation of the VL domain, yielding enthalpically favorable interactions.
3. A network of hydrophilic and hydrophobic interactions, including water-mediated hydrogen bonds, CH–π and CH–O interactions, and aromatic stacking that contribute to high-affinity binding.
4. Distinct water-mediated interaction patterns among PG analogues, reflecting their subtle chemical differences. In particular, the absence of the C5–C6 double bond in PGE□ introduces greater flexibility in the α-chain, disrupting C1-carboxyl interactions with key aromatic residues and altering the water channel observed in PGE□ and ONO5 complexes. These changes correlate with reduced conformational rigidity and lower overall crystallographic temperature factors in PGE□-bound structures.
5. A binding crevasse enriched in lateral hydrogen-bonded chains along the ceiling of the cavity, forming a structurally favorable environment for hydrophobic PG moieties with minimal dehydration penalty.

**Figure 1.**
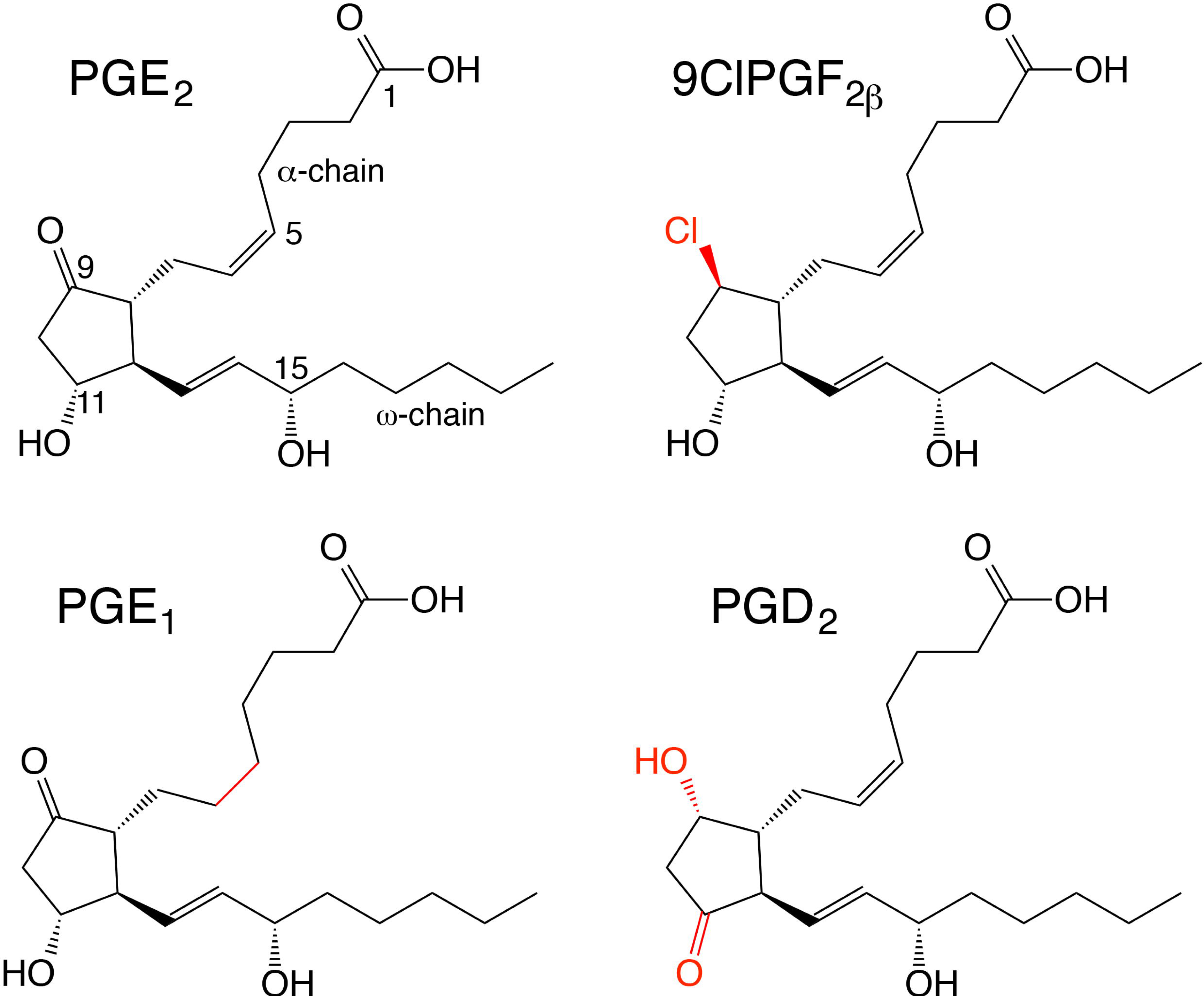
Chemical structures of prostaglandins used in this study. Chemical structures of PGE□, PGE□, PGD□, and ONO5 (9-chloro-PGF□β). Structural features that differ from PGE□ are highlighted in red.

Thermodynamic analyses revealed that PG binding to the Fab is strongly enthalpy-driven. While the intact mAb exhibits high specificity among PG analogues—consistent with ELISA data—such differences are less apparent in Fab binding under dilute solution conditions. Notably, ITC profiles differed between Fab and mAb: Fab titrations fit well to a simple one-site binding model, whereas mAb titrations displayed systematic deviations, suggesting antigen-binding affinity changes during titration or contributions from interdomain coupling within the intact immunoglobulin. To probe these effects further, we examined PG binding under macromolecular crowding conditions using high concentrations of bovine serum albumin (BSA). In the presence of BSA, the mAb—but not the Fab—exhibited markedly improved discrimination between PGE□ and PGE□, reflecting diminished entropy–enthalpy compensation and stabilization of domain–domain interactions in the crowded milieu. These results parallel observations from ELISA systems, where crowding at the solid–liquid interface enhances antigen specificity (11). Collectively, these findings provide mechanistic insight into the structural and thermodynamic principles governing high-fidelity recognition of chemically similar lipid ligands by antibodies and underscore the importance of bound-water architecture and macromolecular environment in shaping antigen discrimination.

## Results

### 3.1. Crystal structures of the Fab in the antigen-free state and in complexes with PGE□, PGE□, and ONO5

We determined crystal structures of the anti-PGE□ Fab fragment in its antigen-free form and in complex with PGE□, PGE□, or ONO5 at resolutions of 2.0 Å, 1.9 Å, 1.7 Å, and 1.7 Å, respectively (Table 1). Fab samples were prepared using an optimized purification procedure, yielding highly homogeneous material suitable for crystallographic analysis by twice yield with adding 2-propanol in the final elution buffer of gel filtration chromatography, as described under Experimental Procedures and previously reported (Supporting Appendix) (13).

**Table 1.** Statistics of the collected diffraction data and the refined structural model.

|  | Fab | Fab: PGE<sub>2</sub> | Fab: PGE<sub>1</sub> | Fab: ONO5 |
| --- | --- | --- | --- | --- |
| A. Data collection |  |  |  |  |
| Space group | *P* 2<sub>1</sub>2<sub>1</sub>2 | *P* 2<sub>1</sub>2<sub>1</sub>2<sub>1</sub> | *P* 2<sub>1</sub>2<sub>1</sub>2<sub>1</sub> | *P* 2<sub>1</sub>2<sub>1</sub>2<sub>1</sub> |
| Cell dimensions *a*, *b*, *c* (Å) | 90.89, 141.53, 40.26 | 70.26, 82.53, 82.74 | 70.46, 82.03, 82.31 | 70.66, 82.31, 82.56 |
| Resolution (Å) | 2.02 (2.09-2.02)<sup>*</sup> | 1.90 (1.97-1.90) | 1.70 (1.79-1.70) | 1.70 (1.79-1.70) |
| Total observations | 253553 (36623) | 280174 (40116) | 389826 (55653) | 370452 (35445) |
| Unique reflections<sup>a)</sup> | 34946 (3405) | 38574 (3795) | 53130 (5241) | 53167 (4889) |
| R<sub>merge</sub> (%) | 9.7 (39.6) | 13.8 (40.6) | 8.4 (40.3) | 10.0 (40.6) |
| Mean *I* / $\sigma$ (*I*) | 11.9 (4.4) | 9.7 (4.4) | 14.9 (4.7) | 15.8 (3.8) |
| Completeness (%) | 99.9 (99.8) | 100.0 (100.0) | 100.0 (99.9) | 99.2 (94.4) |
| B. Refinement statistics |  |  |  |  |
| R<sub>factor</sub> / R<sub>free</sub><sup>a)</sup> (%) | 18.6/21.9 (23.6/27.5) | 16.7/19.6 (20.1/25.1) | 16.7 / 18.8 (20.1/24.1) | 17.2/18.9 (23.5/25.1) |
| Protein atoms | 3277 | 3301 | 3334 | 3310 |
| Ligand atoms | 5 | 25 | 25 | 25 |
| Water molecules | 531 | 572 | 726 | 711 |
| Average B-factors |  |  |  |  |
| Protein (Å<sup>2</sup>) | 29.3 | 22.0 | 19.3 | 20.9 |
| Ligand (Å<sup>2</sup>) | 43.5 | 17.0 | 16.4 | 13.3 |
| Water molecules (Å<sup>2</sup>) | 39.1 | 33.1 | 32.1 | 31.6 |
| R.m.s deviation from ideal |  |  |  |  |
| Bond length(Å) | 0.007 | 0.005 | 0.006 | 0.005 |
| Bond angle (deg.) | 1.11 | 1.07 | 1.06 | 1.02 |
| Ramachandran plot (%) |  |  |  |  |
| Favored (Allowed) region | 94.7 (4.3) | 96.2 (3.3) | 96.6 (2.9) | 96.5 (3.2) |
| Outliers region | 1.0 | 0.5 | 0.5 | 0.3 |
| PDB ID | 3WE6 | 3WFH | 3WHX | 3WIF |

<sup>\*</sup>) The values in the parentheses refer to the highest resolution shell.
<sup>a)</sup> Ten percent of the reflections were set aside for R<sub>free</sub> calculation.

The mouse IgG1κ antibody comprises six immunoglobulin-fold domains: two Fab arms containing the VH and VL variable domains and the CH1 and CL constant domains, and an Fc region composed of CH2 and CH3 domains. The Fab structure solved here corresponds to the VH, VL, CH1, and CL domains, consistent with the cDNA sequences and partial N-terminal and trypsin-digested fragment sequencing of the papain-digested mAb fragments (14, 15).

#### Overall structural comparison

The antigen-free Fab structure contains a bound 2-propanol molecule, originating from the gel-filtration buffer used in the final purification step. This molecule occupies the antigen-binding crevasse in a position corresponding to the C11 hydroxyl region of PG, and is supported by strong residual electron density. In contrast, the PGE□, PGE□, and ONO5 complex structures show well-defined density for each ligand fully occupying the binding crevasse (Fig. 1, 2).

Superposition of the antigen-free Fab with any PG-bound structure reveals a substantial conformational difference, with rms deviations of >0.8 Å across all atoms (Table 1). By contrast, the PG complexes are highly similar to one another, with rms deviations <0.3 Å. When each domain (VH, VL, CH1, CL) is superimposed independently, rms deviations remain <0.3 Å, confirming that antigen binding induces coordinated but localized structural adjustments.

#### Elbow angle differences

The Fab elbow angle—the angle between variable and constant domains—differs significantly between antigen-free and ligand-bound states. The antigen-free Fab exhibits an elbow angle of 183.1° ± 0.6°, whereas all PG-bound complexes share an increased elbow angle of 186.4° ± 0.6°. This consistent shift across all complexes indicates that PG binding induces a reproducible relative domain rotation.

#### Architecture of the antigen-binding crevasse

The Fab possesses a deep, ∼15-Å crevasse at the VH–VL interface. This crevasse contains two solvent-accessible openings:

1. A wide “mouth” at the C1-carboxyl end of the ligand, corresponding to the site used for antigen conjugation in hybridoma screening.
2. A narrower water channel, running from the C11 and C15 hydroxyl region toward bulk solvent. This channel is present in the antigen-free Fab and in the PGE□ complex but becomes altered or occluded in other PG complexes (Fig. 1, 2).

#### Ligand conformations and interactions

In the Fab:PGE□ complex, the bound PGE□ is fully ordered and its conformation closely resembles that of crystallized free PGE□ (CCDC:1238093), except for modest differences in the flexible α- and ω-chain termini (Fig. 2B). A total of 16 amino acid residues form the binding environment, engaging both hydrophilic and hydrophobic interactions, many mediated by buried water molecules (Fig. 2, 3).

**Figure 2.**
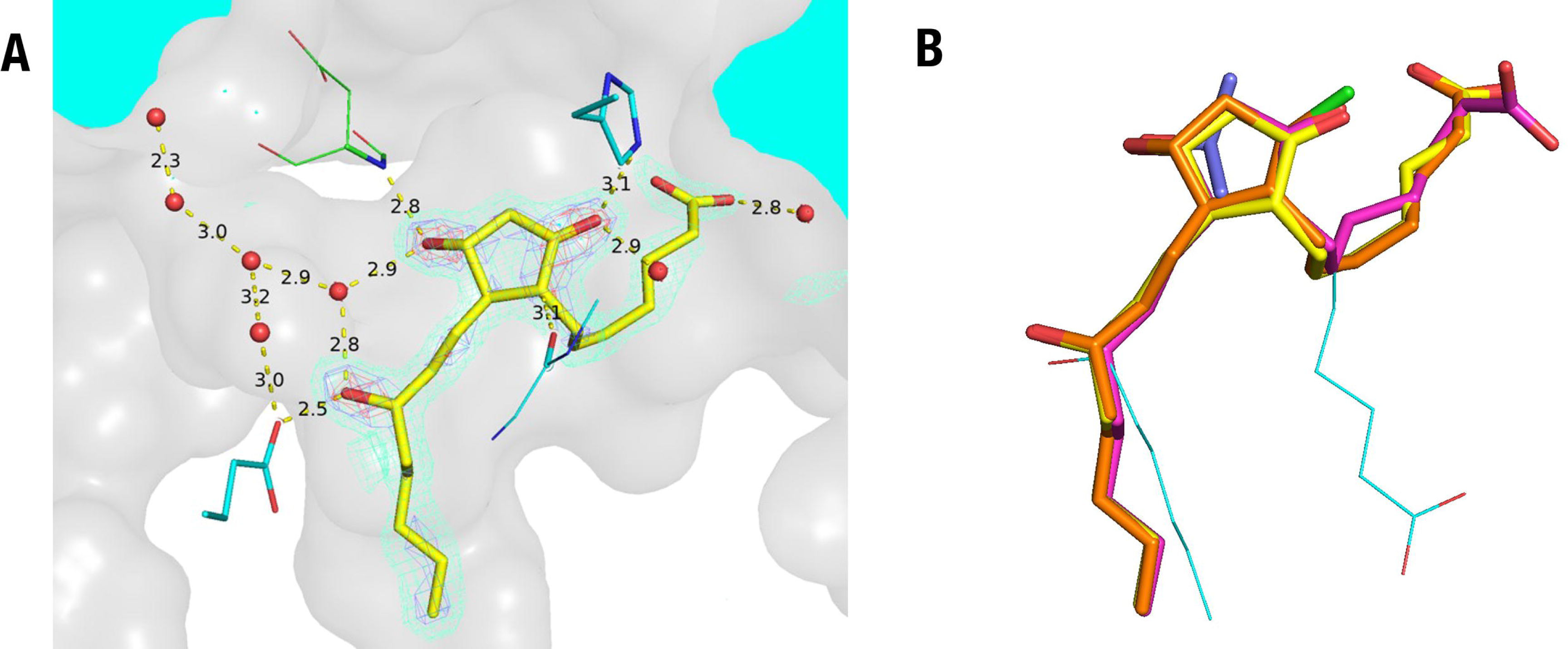
Structural features of bound prostaglandins in Fab complexes. **A.** Bound PGE□ within the Fab binding crevasse, showing direct polar interactions and water-mediated hydrogen bonds. Key water molecules (w1, w2, w3) are indicated with hydrogen-bond distances (Å). The PGE□ omit electron density map is shown contoured at σ = 3 (sky-blue mesh) and σ = 5 (red). VH residues are shown in stick representation (green: C; red: O; blue: N), VL residues in sky-blue, and PGE□ in yellow. Protein surface and solvent region are shown in gray and Sky blue, respectively. **B.** Superposition of bound PGE□ (yellow), PGE□ (pink), ONO5 (orange; chloride in green), and 2-propanol from the antigen-free Fab structure (purple). The conformation of crystallized PGE□ (CCDC 1238093) is shown in thin sky-blue.

**Figure 3.**
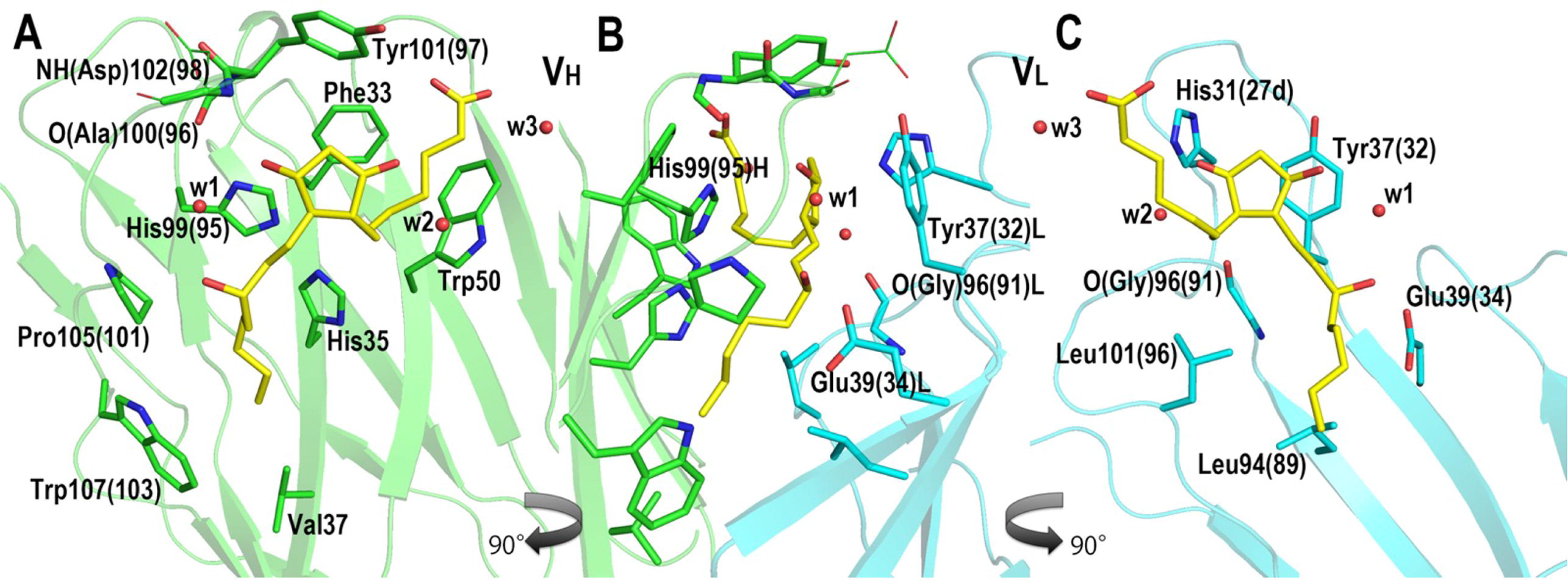
Polar and hydrophobic interactions in the PGE□:Fab complex. **A.** Interactions between PGE□ and residues of the VH domain. **B.** Same view rotated 90°, showing interactions with both VH and VL domains. **C.** View rotated 180°, highlighting interactions between PGE□ and VL. Atom colors same as Fig. 2A.

The hydrophilic moiety of PGE□ consists of four oxygenated functional groups (one carboxyl, one carbonyl, two hydroxyls). These groups participate in a network of direct and water-mediated hydrogen bonds:

1. C1-carboxyl group interacts with water molecule w3 at the solvent boundary.
2. C9-carbonyl oxygen accepts hydrogen bonds from the His31(27D)L (Kabat numbering in parenesis as the amino acid sequence number, if difference, 14) side chain and water w2.
3. C11-hydroxyl group forms hydrogen bonds with the backbone NH of Asp102(98)H and water w1.
4. The C15-hydroxyl group hydrogen-bonds to Glu39(34)L via w1, completing a water-mediated linkage across the ω-chain.

In addition, the backbone carbonyl of Gly96(91)L is positioned for a CH–O interaction with the C8 carbon of PGE□.

The C1-carboxyl group of PGE□ interacts with three aromatic residues—Phe33H, Trp50H, Tyr101(97)H—via CH–O contacts, stabilizing the α-chain orientation. A continuous water chain emerging from the C11/C15 hydroxyl region extends to solvent, producing a through-hole that is characteristic of PGE□ binding (Fig. 2).

#### Hydrophobic interactions

The cyclopentane ring of PGE□ is nestled among several aromatic and aliphatic residues. Notably, Tyr37(32)L forms a CH–π interaction with the C11 hydrogen at ∼3.7 Å. Additional residues—including Phe33H, His35H, Trp50H, His99(95)H, Tyr101(97)H, Trp107(103)H, Val37H, Pro105(101)H, Leu94(89)L, and Leu101(96)L—lie within 4 Å of PGE□, forming an extensive hydrophobic enclosure.

#### Comparison among PG analogues

Although the binding modes of PGE□, PGE□, and ONO5 share central features, several subtle but functionally important distinctions emerge. In Fab:PGE□, the absence of the C5–C6 double bond increases α-chain flexibility, leading to loss of CH–O interactions with Phe33H, Trp50H, and Tyr101(97)H. Water w3/w3’’ is replaced by two alternative waters (w3’□ and w3’□). The altered α-chain geometry narrows and partially closes the secondary water channel (Fig. 4B), consistent with modified conformational dynamics and lower crystallographic B-factors than the other PG complexes (Fig. 6B).

**Figure 4.**
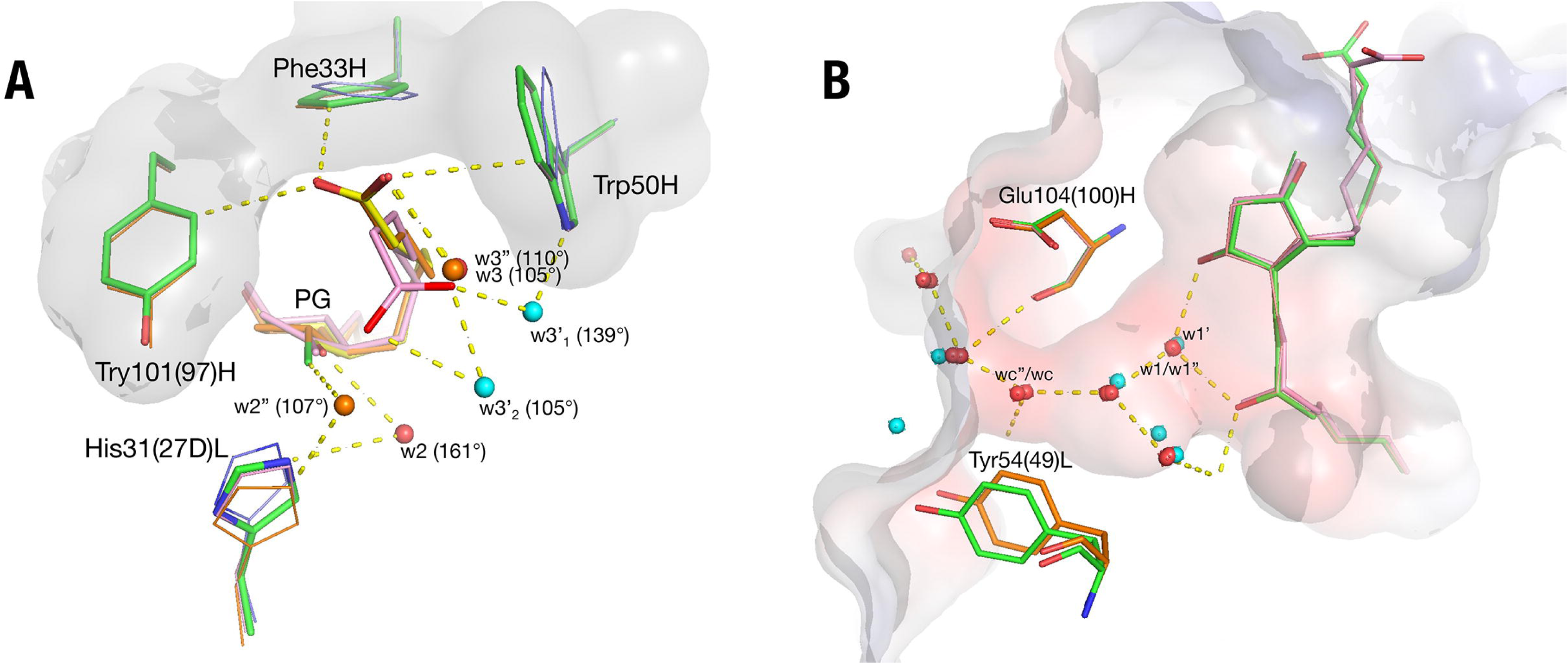
Comparison of water networks and **α**-chain conformations among PG-bound Fab structures. **A.** Structural differences in α-chain interactions and water networks for PGE□ (green), ONO5 (thin orange), and PGE□ (thick pink). Water molecules w2 and w3 (PGE□), w2″ and w3″ (ONO5), and w3′□ / w3′□ (PGE□) are shown. **B.** Closure of the water channel in the PGE□ complex due to repositioning of Tyr54L and Glu104(100)H and subtle shifts in the CDR-H3 loop. Corresponding waters (w1, w1″, w1′) are indicated for PGE□, ONO5, and PGE□ structures.

In Fab:ONO5, the β-chloro substituent at C9 forms a hydrogen bond with His31(27D)L and induces repositioning of water w2→w2’’, and this rearrangement preserves overall binding geometry but alters local hydrogen-bond angles (161° vs. 107° in PGE□) with w2 and w2”, respectively.

Across all structures, aromatic residues maintain face-to-face or edge-to-face orientations relative to PG rings, creating a robust CH–π and π-stacking landscape with >20 Å² buried area per aromatic side chain.

### 3.2. Hydrogen-bonded cage of the crevasse interior

The interior of the antigen-binding crevasse is reinforced by an extensive lateral hydrogen-bonded network formed by side chains and backbone groups of residues lining the VH and VL domains. This hydrogen-bonded “cage” is present not only in the PG-bound structures but also in the antigen-free Fab, although its geometry and hydration patterns differ depending on the ligand bound.

Three predominant hydrogen-bonded chains contribute to the structural integrity and stabilization of the binding crevasse (Supplementary Fig. 2) ;

A: A chain spanning the VH β-sheet scaffold. This network includes Trp47H, His35H, His99(95)H, and the backbone carbonyl of Ala100(96)H along the five-stranded antiparallel β-sheet of the VH domain (Supplementary Fig. 2). These interactions rigidify the wall of the cavity against which the cyclopentane ring of PG packs.
B: A chain running along the ω-chain region of the bound ligand. This includes Trp107(103)H, Tyr41H, Glu39(34)L, a bridging water molecule, and the backbone NH of Lys55(50)L. These form a continuous hydrogen-bonded path parallel to the ω-chain of PGE□, stabilizing the ligand’s orientation within the cleft.
C: A cluster surrounding the large opening of the crevasse. Lys55L, Asp102(98)H, Tyr37(32)L, Tyr101(97)H, and two conserved structural waters create a hydrogen-bonded array capping the cyclopentane ring region at the entrance of the cleft. Together, these networks of hydrogen-bonded side chains and β-sheet backbone groups generate a topologically consistent cage that encases the ligand between the VH and VL β-sheets (Fig. 2, 3; Supplementary Fig. 2). The hydrophobic interior—lined primarily with aromatic residues—provides an apolar environment complementary to the PG hydrocarbon framework, while the hydrogen-bonded networks reinforce the overall architecture of the crevasse.

Importantly, the enrichment of aromatic residues in this interior surface, many of which are involved in hydrogen-bonding networks or π-stacking interactions, limits the number of available polar donors/acceptors. Thus, hydration patterns within the crevasse are highly selective and structurally significant. The buried waters that remain play essential roles in mediating PG recognition, bridging interactions between ligand and protein, and shaping the cavity for optimal complementarity to each PG analogue.

These hydrogen-bonded chains collectively ensure cavity stability and contribute to the enthalpic signature of PG complex formation, consistent with the thermodynamic data presented below.

### 3.3. Conformational changes induced by ligand binding

Prominent conformational changes occur upon prostaglandin binding, most notably within the CDR-H1 loop of the VH domain and in the relative orientation of the Fab variable and constant domains. These structural adjustments are accompanied by reorganization of bound water molecules and are consistent across all PG complex structures.

#### Formation of a short 3□□ helix in CDR-H1

In the antigen-free Fab, the CDR-H1 loop adopts an extended conformation, and the phenyl side chain of Phe29H projects outward toward solvent. The electron density for this side chain is weak, reflecting substantial flexibility (Fig. 5A). Upon binding any of the PG ligands, Phe29H flips inward into a hydrophobic pocket formed between the CDR-H1 and CDR-H2 loops, initiating a cooperative rearrangement that includes: formation of a short 3□□ helix within the CDR-H1 loop, burial of Phe29H within the inter–β-sheet cavity, extrusion of the Glu31H side chain toward solvent, and restructuring of the aromatic cluster formed by Tyr27H and Tyr32H, which engage in π–π stacking interactions (Fig. 5B and 5C).

**Figure 5.**
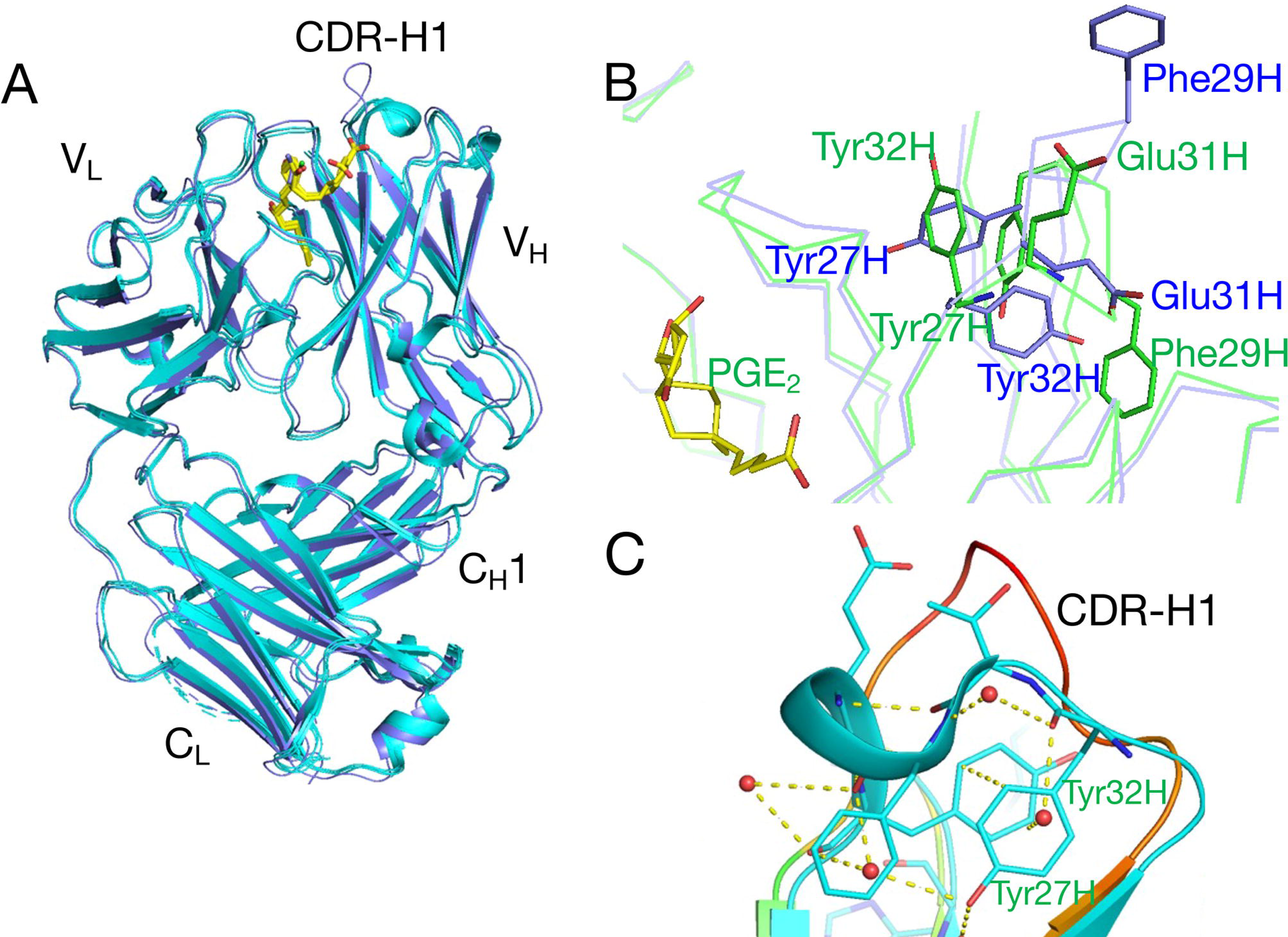
Ligand-induced conformational changes in CDR-H1. **A.** Superposition of antigen-free Fab (purple) and PG-bound Fabs (sky blue) showing overall loop repositioning. **B.** Detailed comparison of the CDR-H1 loop between antigen-free (purple) and PGE□-bound (green) structures, highlighting Tyr27, Tyr32, Phe29, and Glu31 side chain rearrangements. **C.** Hydrogen-bond network stabilizing the short 3□□ helix in the PGE□-bound CDR-H1 loop, including CH–O and π–π interactions and key water molecules, superposed with the antigen-free loop (orange).

This helix formation is stabilized by multiple hydrogen bonds and “capping” water molecules, which help lock the loop into its ligand-bound conformation. Collectively, these changes result in a significant reduction of exposed hydrophobic surface area and contribute favorably to the enthalpic component of binding, consistent with calorimetric measurements.

#### Domain-level motions

Ligand binding induces small but measurable adjustments in the orientation of the Fab domains: The VL domain undergoes an approximately 1° rotation relative to its position in the antigen-free Fab (Supplementary Fig. 3). The elbow angle between the V and C domains increases uniformly across all PG-bound structures, as noted earlier. These domain motions, although subtle, are highly reproducible among the three ligand-bound structures and reflect the mechanical coupling of antigen recognition to global Fab architecture.

#### Changes in temperature factors and dynamic features

Analysis of crystallographic temperature (B) factors reveals distinct patterns of dynamic behavior: In the antigen-free Fab, the CDR-H1 loop of the VH domain displays the highest B-factors, consistent with its flexibility and absence of stabilizing interactions (Fig. 6A). In the PG-bound structures, the CDR-H1 loop becomes more ordered due to helix formation and aromatic packing, whereas a different region—the antipodal loops of the CL domain—exhibits markedly elevated B-factors relative to the antigen-free structure (Fig. 6).

**Figure 6.**
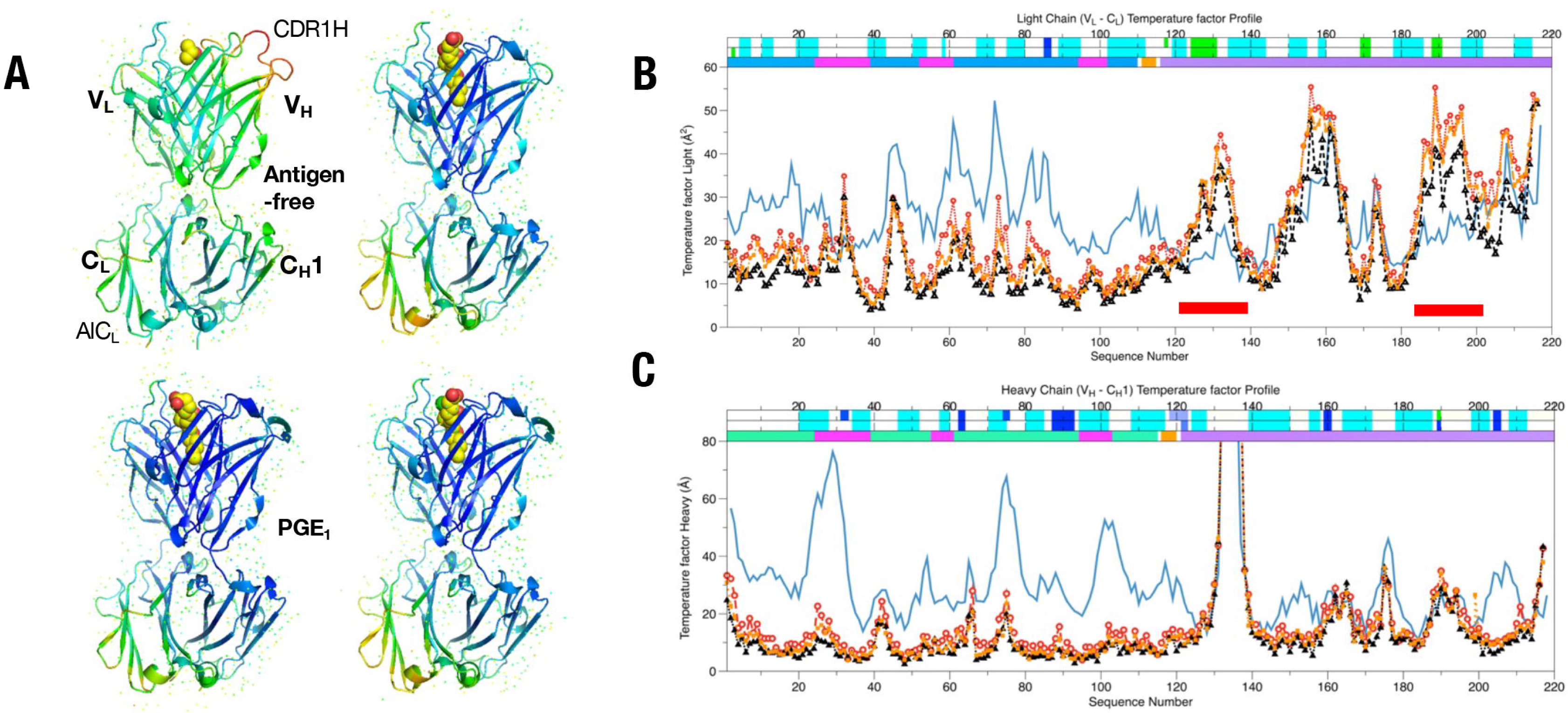
Domain-level thermal motion and temperature-factor distributions. **A**. Crystallographic B-factor distribution mapped onto Fab structures comparing antigen-free and PG-bound Fab structures. **B**. Increased B-factors in the CL domain loops of PG-bound Fabs versus the antigen-free Fab, contrasted with decreased B-factors in the VH CDR-H1 loop upon ligand binding. These CL loop regions correspond spatially to the expected CH2–CL interface of intact IgG (PDB 1IGY). Upper: Light chain (V_L_ and C_L_). Lower: Heavy chain (V_H_, and C_H_1). B-factor of antigen free Fab (blue line, ——). Fab:PGE_2_ (red line, —○—), Fab:PGE_1_ (black line, —△—), Fab:ONO5 (orange line,—□—); Bar legend: Secondary structures of complexes (Upper bar) and antigen free (Middle bar), β-sheet (sky blue) / Helix(blue). Bottom bar: domain of V_H_ (green) / V_L_ (blue) with CDR (pink) and C_H_1 / C_L_ (violet). Antipode CL domain loops (thick red lines).

The increased mobility of the CL loops in the PG-bound state is particularly notable for PGE□:Fab, which displays slightly lower B-factors in these loops than the PGE□ and ONO5 complexes within the same crystal lattice. This result is consistent with subtle structural mismatches introduced by PGE□’s more flexible α-chain.

Superposition with an intact IgG1 crystal structure (PDB 1IGY) (18) suggests that the CL loops showing increased mobility reside near the expected interface with the CH2 domain of the Fc region. Thus, ligand-induced changes in the Fab variable region appear to propagate into the constant domains, influencing overall Fab/Fc dynamics. This coupling may contribute to the distinct thermodynamic behavior observed for the intact mAb compared with the isolated Fab, as described below.

### 3.4. Thermodynamics of PG binding to the Fab

To elucidate the thermodynamic properties underlying antigen recognition by the Fab fragment, we performed both isothermal titration calorimetry (ITC) and differential scanning calorimetry (DSC). These measurements reveal that PG binding to the Fab is uniformly enthalpy-driven, consistent with the extensive network of hydrogen bonds, aromatic interactions, and water-mediated contacts observed crystallographically.

#### ITC analysis of Fab–PG interactions

Binding of PGE□ to the Fab was strongly exothermic, with a measured binding enthalpy Δ*H*_b_ of –14.8 ± 0.4 kcal·mol□¹ (–61.9 kJ·mol□¹) and the dissociation constant was *K*_d_ = 23 ± 123 nM with one site fitting model, and the entropy was deduced enthalpy to be –TΔ*S*_b_ = 4.4 ± 0.3 kcal·mol□¹ at 25 °C (298 K) as Δ*G*_b_ = –10.4 ± 0.1 kcal·mol□¹ (n=3), respectively (Table 2 and Supplementary Fig. 1A). These values confirm that the binding is primarily driven by enthalpic contributions, with entropy contributing moderately and unfavorably—consistent with the conformational ordering of CDR loops and internal water molecules.

**Table 2.**
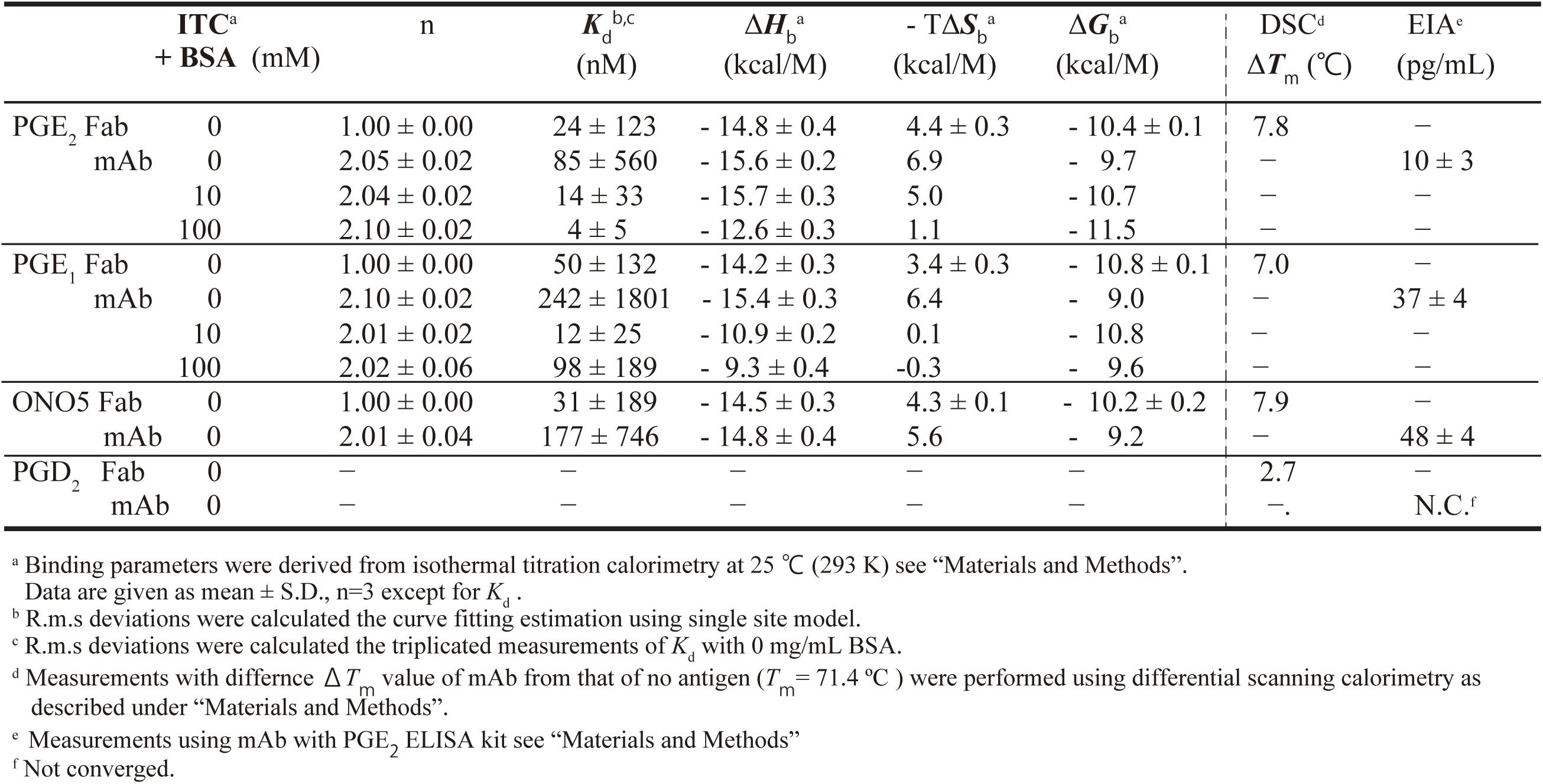
Binding parameters of Anti-PGE2 mAb and Fab.

Importantly, the calculated free energies (Δ*G*_b_) for PGE□, PGE□, and ONO5 were similar under these dilute-solution ITC conditions, despite the clear differences detected by enzyme immunoassays (EIA) (Supplementary Fig. 5) (11). This suggests that the Fab fragment alone does not capture the full extent of discriminatory capacity observed in the intact mAb.

#### DSC analysis of Fab stability in complex with PGs

Thermal unfolding of the Fab in the presence and absence of ligand was monitored by DSC. The antigen-free Fab exhibited a melting temperature (*T*_m_) of 71.4 °C (Table 2, Supplementary Fig. 1B). Binding of any of the PG ligands increased the thermal stability of the Fab, reflected in positive shifts in *T*_m_; Differences of *T*_m_ from antigen free Δ*T*_m_ for PGE□, PGE□, ONO5 and PGD□ were 7.8 °C, 7.0 °C, 7.9 °C and 2.7 °C, respectively. Although we were unable to crystallize a PGD□ complex, its intermediate stabilizing effect is consistent with its closer structural similarity to PGE□ relative to other prostanoids (Fig. 1). The thermal unfolding profiles for PGE□, PGE□, and ONO5 complexes superimposed well, and their excess transition enthalpies were comparable, indicating similar global stabilization mechanisms across the three complexes.

The slightly lower Δ*T*_m_ for PGE□ aligns with the structural observations that its flexible, single-bonded C5–C6 region disrupts optimal packing and water-mediated interactions relative to PGE□ and ONO5.

Together, the ITC and DSC data demonstrate that ligand binding significantly stabilizes the Fab and that the recognition process is uniformly enthalpy-driven, consistent with the extensive hydrogen bonding and ordered water networks identified in the high-resolution crystal structures.

### 3.5. PG binding to the intact mAb in ITC under dilute and crowded conditions

To compare the thermodynamic behavior of ligand binding to the intact monoclonal antibody (mAb) with that of the isolated Fab, we performed ITC experiments with the full-length anti-PGE□ IgG1κ. These measurements were further evaluated under macromolecular crowding conditions using bovine serum albumin (BSA), in order to approximate the high-protein environment of blood plasma. This approach aimed to capture the interdomain coupling effects between the Fab and Fc regions that are absent when studying the Fab fragment alone.

#### Differences between mAb and Fab in ITC binding profiles

In contrast to the Fab, ITC titrations of the intact mAb exhibited; greater overall exothermicity (more negative Δ*H*_b_), larger unfavorable entropic contributions (–TΔ*S*_b_), and more negative binding free energies (Δ*G*_b_), indicating higher apparent affinity for PGE□, PGE□, and ONO5 compared with the Fab fragment (Figure 7, Table 2). However, the titration curves for the mAb displayed systematic deviations from a simple one-site binding model, regardless of the titration by PGE□ or PGE□. These deviations suggest that antigen binding to the intact antibody may involve; (1) Non-equivalent binding behavior at the two Fab arms. (2) Progressive tightening of affinity during titration. or (3) Dynamic allosteric coupling between Fab and Fc domains.

**Figure 7.**
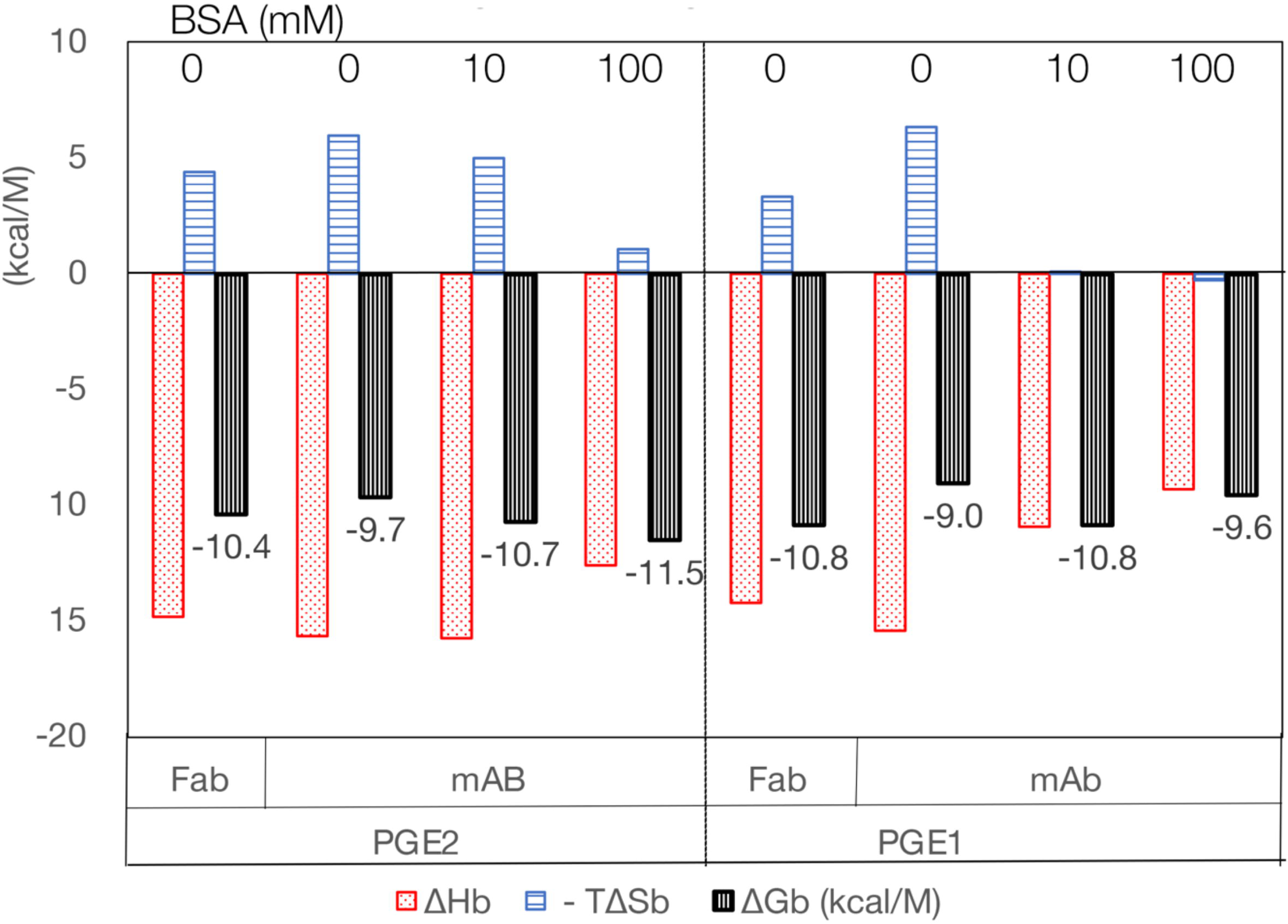
Thermodynamic parameters for PG binding to Fab and mAb. ITC-derived binding enthalpy (Δ*H*_b_), entropy term (–TΔ*S*_b_), and free energy (Δ*G*_b_) for PGE□ and PGE□ binding to the Fab and the intact mAb in the presence or absence of BSA (macromolecular crowding). Results illustrate enthalpy-driven binding, entropy–enthalpy compensation, and enhanced ligand discrimination under crowded conditions.

Forcing a one-site model fit minimized overfitting but produced unusually large residuals, consistent with structural and dynamic complexity not present in the isolated Fab (Supplementary Fig. 4).

#### Enhanced discrimination between PGE□ and PGE□ under crowded conditions

To further probe differential ligand recognition, ITC experiments were repeated in the presence of 10–100 mg/mL BSA, representing a physiologically relevant crowding environment. Strikingly, under these crowded conditions (Fig. 7 and Table 2).

The intact mAb—but not the Fab—showed clear thermodynamic discrimination between PGE□ and PGE□, in agreement with EIA results (Table 2 and Supplementary Fig. 5). For PGE□ binding, crowding caused a larger increase in entropic penalties (–TΔ*S*_b_) than in enthalpic loss (Δ*H*_b_), yielding more negative Δ*G*_b_ and thus higher affinity. For PGE□ binding, crowding increased enthalpic loss more than entropic penalty, resulting in less favorable Δ*G*_b_ relative to PGE□.

This pattern indicates that the intact mAb experiences reduced entropy–enthalpy compensation under crowding, a known phenomenon in macromolecular assemblies. Such conditions magnify subtle structural mismatches—such as the flexible α-chain conformation of PGE□—leading to enhanced discrimination between closely related prostanoids.

#### Simplification of mAb titration behavior in the presence of BSA

The addition of BSA dramatically improved the quality of single-site model fitting for both PGE□ and PGE□ titrations. Fit residuals were greatly reduced, Binding curves became more symmetrical, and replicate titrations were more reproducible (Table 2; Supplementary Fig. 4).

These improvements suggest that crowding effectively decouples the two antigen-binding sites or stabilizes the mAb in a conformation with more uniform binding behavior. This is consistent with the idea that high-protein environments restrict domain motion and dampen the structural heterogeneity responsible for fitting deviations under dilute conditions.

#### Consistency with EIA and crystallographic observations

The enhanced specificity for PGE□ over PGE□ observed for the intact mAb under crowding conditions mirrors; the superior specificity measured by EIA (Supplementary Fig. 5) (11), and the structural distinction observed between PGE□:Fab and PGE□/ONO5:Fab, particularly the loss of C1–aromatic interactions and collapse of the water channel in the PGE□ complex (Fig. 4–6), in consistent with those of DSC Δ*T*_m_ (Supplementary Fig. 1B and Table 2).

Together, these data support the conclusion that macromolecular crowding amplifies subtle structural incompatibilities and modulates domain dynamics, thereby enhancing the intrinsic discrimination capacity of the monoclonal antibody.

## Discussion

In this study, we elucidated the molecular principles governing the highly specific recognition of PGE□ by an anti-PGE□ monoclonal antibody using high-resolution crystal structures of Fab–ligand complexes and comprehensive thermodynamic analyses. By comparing PGE□, PGE□, and ONO5, we were able to dissect the structural determinants and water-network architectures that discriminate among closely related prostanoid ligands. The results underscore the essential role of ligand-induced conformational changes, water-mediated interactions, and macromolecular environmental factors in shaping antigen specificity.

### Structural basis of ligand discrimination

The high-resolution Fab structures revealed a pronounced conformational transition between the antigen-free state and the PG-bound states. These changes include the formation of a short 3□□ helix in CDR-H1, burial of Phe29H, rearrangement of aromatic stacking interactions, and reorganization of structured water networks. Such coordinated structural reconfigurations indicate that PGE□ binding is accompanied by a dynamic recognition mechanism rather than a purely lock-and-key interaction.

The distinct structural features of the PG analogues—particularly the flexible α-chain of PGE□ lacking the C5–C6 double bond—produce measurable differences in how each ligand fits into the Fab crevasse. Loss of optimal CH–O contacts and altered water-mediated interactions for PGE□ correspond with the closure of a secondary water channel and subtle shifts in the positions of key residues on the CDR-H3 and VL loops. These differences align with the lower crystallographic B-factors in Fab:PGE□ compared with Fab:PGE□ and Fab:ONO5, suggesting altered local dynamics consistent with suboptimal complementarity.

The present study highlights the essential contribution of bound water molecules in ligand discrimination. Specific water positions differ between PG complexes, particularly near C1 and C11–C15 hydroxyl groups, forming bridges critical for anchoring the ligand. Experimental structural insights of this nature are currently beyond the predictive capability of AI-based modeling tools such as AlphaFold 3, which, despite major advances in multimer and ligand prediction accuracy (19), do not yet incorporate explicit bound water structures. Our findings therefore provide valuable empirical reference points for understanding water-mediated specificity, and water network and helix structure could be contributed the dynamic and precise recognition of ligand while β-sheets as stable scaffold thermodynamically (Fig. 4 to 6).

### Implications for ligand design and receptor recognition

The differential recognition of PGE□ and PGE□ by this antibody parallels their distinct pharmacological effects at EP receptors (6, 10, 22). Subtle chemical variations in prostanoids—such as double-bond geometry, chain flexibility, and functional group placement—translate into meaningful differences in how water networks and aromatic interactions stabilize ligand binding. These results highlight core principles applicable to antibody engineering, receptor-ligand studies, and structure-guided drug design. In particular, the present work underscores that enthalpy–entropy compensation in biomolecular recognition (23) is strongly modulated by the conformational and hydration landscape within binding pockets. Engineering strategies that optimize water networks or alter loop dynamics may therefore offer powerful leverage in tuning specificity and affinity in future antibody or ligand design efforts (20, 21).

### Thermodynamic and biophysical consequences of macromolecular crowding

Strikingly, while the isolated Fab fragment displayed similar binding free energies (Δ*G*_b_) toward PGE□, PGE□, and ONO5 in dilute ITC conditions, the intact mAb exhibited substantial differences in affinity that became even more pronounced under macromolecular crowding. In crowded environments containing high concentrations of BSA, the mAb displayed clearly enhanced discrimination between PGE□ and PGE□, consistent with the specificity observed in ELISA assays (Supplementary Fig. 5) (11). This enhanced discrimination arises from diminished enthalpy–entropy compensation under crowded conditions, where entropy contributions associated with water rearrangement or domain flexibility become suppressed. Crowding effectively limits the conformational heterogeneity of the intact antibody, stabilizing domain-domain interactions and amplifying subtle structural differences in PG binding. The simplification of mAb ITC profiles and improved fitting to a single-site binding model in the presence of BSA strongly supports this interpretation.

These observations highlight the functional relevance of studying antibody recognition in physiologically relevant conditions, especially given that the Fc domain contributes to paratope stabilization and epitope specificity (24). The propagation of ligand-induced conformational changes from the Fab to the constant domains—reflected in altered B-factor patterns—further supports the interconnected nature of antibody architecture.

### Conclusions

Together, our structural and thermodynamic analyses provide a comprehensive mechanistic framework for understanding how the anti-PGE□ mAb achieves high-fidelity discrimination of closely related prostanoid ligands. The work emphasizes:

- the key role of bound water networks in antigen recognition,
- ligand-induced conformational changes that optimize enthalpic contributions,
- the importance of aromatic and CH–O/CH–π interactions,
- structural differences among PG analogues that translate into measurable thermodynamic consequences, and
- the significant influence of the macromolecular environment on antibody specificity.

These findings not only advance our understanding of prostanoid recognition but also provide generalizable principles applicable to antibody engineering, ligand optimization, and the design of next-generation therapeutics targeting lipid mediators.

## Experimental Procedures

### Preparation of Fab Fragment

The anti-PGE□ monoclonal antibody mAb and its Fab fragment as a mouse IgG1κ were produced from serum-free hybridoma culture supernatants as previously described (11, 13), with modifications to improve Fab yield and purity with cation, anion and gel filtration chromatography (see Supporting Appendix). Following mAb purification by proteinA affinity chromatography and papain digestion was performed to generate Fab fragments (left column of Supporting Appendix 1’) (13).

### Crystallization of Fab

Purified Fab was concentrated to 15 mg/mL using a centrifugal concentrator in the gel-filtration buffer. For crystallization screening, equimolar mixtures of Fab with twofold molar excess of PGE□, PGE□ (Cayman Chemical), ONO5 (Ono Pharmaceuticals), or Fab alone (antigen-free) were prepared. Crystallization trials were conducted at 20 °C using oil-microbatch robotic setups (Douglas Instruments; TERA system at RIKEN) with commercial screening kits (Hampton Research, Emerald Biostructures) (25, 26).

Crystallization conditions were as follows: Antigen-free Fab: 16% (w/v) PEG 8000, 0.1 M sodium phosphate–citrate (pH 4.2), 0.2 M NaCl, 10 mM praseodymium acetate.

Fab:PGE□ complex: 30% (w/v) PEG 4000, 0.2 M ammonium sulfate, 0.1 M sodium citrate (pH 6.0). Fab:PGE□ complex: 25% (w/v) PEG 3350, 0.1 M Bis-Tris-HCl (pH 6.0). Fab:ONO5 complex: 25% (w/v) PEG 3350, 0.1 M sodium acetate (pH 5.0).

### **X-** ray Data Collection and Processing

Diffraction data were collected at 100 K on beamline BL26B2 at the RIKEN SPring-8 Center (27). Crystals were cryoprotected with 24% (v/v) glycerol before flash cooling. Data for the antigen-free Fab, and the Fab complexes with PGE□, PGE□, and ONO5 were collected to resolutions of 2.0 Å, 1.9 Å, 1.7 Å, and 1.7 Å, respectively. Data were processed with MOSFLM and scaled with SCALA from the CCP4 suite (28). Crystallographic statistics are summarized in Table 1.

### Structure Determination and Refinement

The antigen-free Fab structure was solved by molecular replacement using MOLREP (29) with PDB entry 2DDQ (15) as the search model. Iterative cycles of refinement and model building were performed using REFMAC5 (30), CNS (31), PHENIX (32), and COOT (33). Ligand topology and parameter files were generated using PRODRG2 with manual adjustments (34).

Final R/R_free_ values were: for Antigen-free Fab, Fab:PGE□, Fab:PGE□ and Fab:ONO5 as 18.6% / 21.9%, 16.7% / 19.6%16.7% / 18.8% and 17.2% / 18.9%, respectively.

No Ramachandran outliers were present. Structural validation was performed using CCP4 tools and PHENIX. Coordinates and structure factors were deposited in the PDB under accession codes 3WE6, 3WFH, 3WFX, and 3WIF.

Elbow angles were calculated as described by Stanfield *et al.* (16). Domain shift was calculated with LSQKAB in CCP4 (28) (Supplementary Fig. 3). Structural analyses and figures were with PyMOL (Schrödinger, LLC).

### Isothermal Titration Calorimetry (ITC)

Thermodynamic parameters for PG binding were measured using a MicroCal VP-ITC calorimeter (35). Experiments were performed at 25 °C in 10 mM sodium phosphate (pH 7.0) containing 140 mM NaCl. Fab (8.9 μM) or intact mAb (3.5 μM) was placed in the cell; ligands (PGE□, PGE□, or ONO5) were titrated into the sample. Data were analyzed using MicroCal ORIGIN 5.0 with a one-site binding model. Triplicate experiments were performed for *K*_d_ and Δ*H*_b_ determinations. Binding free energy (Δ*G*_b_) and entropy term (–TΔ*S*_b_) were calculated by:

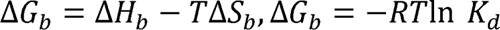

with T = 298 K and R = gas constant.

### Differential Scanning Calorimetry (DSC)

Fab thermal stability was assessed using a VP-capillary DSC (MicroCal) at 200 °C/hour scan rate (36). Fab (0.33 mg/mL; ∼0.1 mM) was mixed with or without each PG ligand (26.7 μg/mL; ∼0.8 mM). Data were analyzed using ORIGIN 7.0. Melting temperatures (*T*_m_) represent averages of triplicate runs.

### Enzyme Immunoassay (EIA)

EIA measurements were performed using a commercial sandwich ELISA kit (Cayman Chemical).

Data analyses were conducted using DataGraph (VisualDataTools) or Prism (GraphPad) as well as Excel (MicroSoft).

## Supporting information

SupportingInfortionTXT

SupportingInfortionfigs

## Data Availability

*Mus musculus* mRNA sequences for the anti-PGE□ mAb heavy and light chains are available in DDBJ under accession numbers AB453396 and AB453397 as an IgG1κ. The ambiguous nucleotides in AB453396 correspond to residues resolved as A and T for the uncertain nucleic acid and the deduced amino acid sequence of W^1353^ and X^435^ in heavy chain, respectively. Coordinates and structure factors for antigen-free Fab and PG-bound complexes had been deposited with PDB accession codes 3WE6, 3WFH, 3WFX, and 3WIF.

## Acknowledgments

We thank Drs. Hitoshi Iino, Akeo Shinkai, Akio Ebihara and Seiki Kuramitsu for allowing our use of ITC equipment, and Drs. Tetsuya Hori and Yasushi Nitanai for valuable comments. We grateful to Dr. Tetsuya Ishikawa for his continuous encourage. ONO5 was kindly gifted by Ono Pharm. The synchrotron radiation experiments were performed at BL26B2 in SPring-8 with the approval of RIKEN.

## Author contributions

MS, MM and SY were experimental design to manuscript writing. HA checked final manuscript. MS, HS, HA, MY and MM for crystallography, MS, MT and MM for ITC, MT and KY for DSC,. MS, ASA and SY were sample preparation and YK and SY for EIA and cDNA sequencings. NT and KT for partial protein sequencing of papain digested mAb eluates of protein G fractions. All experiments and the data verification were responsible for MM. Before finalized the manuscript, KY and SY passed away. The manuscript is dedicated for them.

