## SupportingInfortionTXT for "Structural and Thermodynamic Determinants of Anti-PGE_2_ mAb Specificity"

***Supporting Information***

**Supplementary figures**

Supplementary Fig. 1 Calorimetric Measurements. A: PGE_2_ titration of Fab in ITC. B: DSCs of Fab with (i) PGE_2,_ (ii) ONO5 (iii) PGE_1,_ (iv) PGD_2,_ (v) antigen free. Panel A: ITC of Fab:PGE_2_. Curb fitting was using one site model. Panel B: The *T*_m_ of antigen free mAb (thick blue line) and mAb with PGE_2_ (thin red).

Supplementary Fig. 2 Hydrogen bonding side chains on the PGE_2_ binding crevasse interior. Coloring scheme is the same as Fig. 2. Atomic distances are small character. mAb ITC by Atomic distances are small character. Coloring scheme is the same as Fig. 2.

Supplementary Fig. 3. Rotatory domain-movement of refined structures between PGE_2_ and PGE_1_ complex structures with rotation axis of C_L_ as the significant rotation by 1 degree. See Material and Method.

Supplementary Fig. 4 ITC of PGE_2_ or PGE_1_ titration to mAb with 0 mM, 10mM and 100mM BSA. A: PGE_2_ ITC. B: PGE_1_. BSA conc.: 0 mg/mL (—●—), 10 mg/mL (---△---), 100 mg/mL (…×…).

Curb fitting was with one site binding model.Fitting curves were used single site parameters and the extraordinary large r.m.s. deviation due to systematic deviation, in particular, with no BSA. The presence of BSA achieves more sharpen recognition. Tighter binding mode did not alter the binding mode like as one rigid body of mAb and bound ligand, on the other hand leaky bind becomes more flexible due to damping of BSA collision.

Supplementary Fig. 5 EIA of mAb with PGE_2_, PGE_1_, ONO5, or PGD_2_ competitive inhibition against SOD:PGE_2_ conjugate. PGE_2_ (—●—). PGE_1_ (---○---). ONO5 (---△---). PGD_2_ (…×…).

Supplementary Fig. 6 average of temperature factors <B-factor> of each VH, CH1, VL or CL of crystal structure of Fab:PGE_2_, Fab:PGE_1_ or Fab:ONO5 as well as antigen free Fab.

Supplementary Fig. 7 Secondary Structure assignment with S-S bridge of VH and VL sequences of Fab with Kabat and Chothia Numbering. CDR : complementary determinant region; Sendary Structure: β-Sheet inner sheet (pale Orenge) and outer sheet (Sky Blue), Helix (Blue in White letter); S-S bridge (Brown circle with connected line); Insertion Sequence Number (Red Character on Yellow).

**Appendix: Sample Information.**

Improved Sample Preparation

Papain digested material (left column in Supporting Appendix 1’) was applied to an anion-exchange column (Mono Q 10/100 GL) pre-equilibrated with 25 mM Tris-HCl (pH 8.0) containing 25 mM NaCl. The flow-through fraction containing Fab was subsequently loaded onto a cation-exchange column (Mono S 10/100 GL) equilibrated with 20 mM sodium phosphate (pH 5.5) and eluted with a linear gradient of 20–220 mM NaCl. The major Fab peak eluted at approximately 130 mM NaCl. (Supporting Appendix 1)

This material was further purified by size-exclusion chromatography on a Superdex 200 10/300 GL column equilibrated in 10 mM sodium phosphate (pH 7.0) containing 140 mM NaCl and 5% (v/v) 2-propanol. The purity and homogeneity of the Fab preparation were verified by SDS-PAGE. Partial amino acid sequences of the papain digested mAb were confirmed by Edman degradation and partial sequencing of trypsin-digested Fab and Fc fragments with MS/MS and cDNA sequencing, resolving ambiguities in the nucleotide sequence.

**Appendix Figures**

Supporting Appendix 1: Preparation and characterizations of mAb and Fab.

Supporting Appendix 1’: Conventional Large Preparation of anti-PGE_2_ Fab ^(13).^

Supporting Appendix 2: SDS PAGE and partial amino acid sequences of the purified Papain digested anti-PGE_2_ mouse IgGκ with protein A and G.

Supporting Appendix 3: Anti-PGE_2_ immunoglobin cDNA and deduced amino acid sequence with partial amino acid sequence of Heavy chain of papain digested fragments as Fab and Fc. (Fraction 1 and Fraction2).

Supporting Appendix 4: Sequence alignment of VL and VH in anti-PGE_2_ mAb and related mAbs.
