## SupportingInfortionfigs for "Structural and Thermodynamic Determinants of Anti-PGE_2_ mAb Specificity"

### Supplementary Fig. 1 Calorimetric Measurements

A. Isothermal titration calorimetry（ITC）

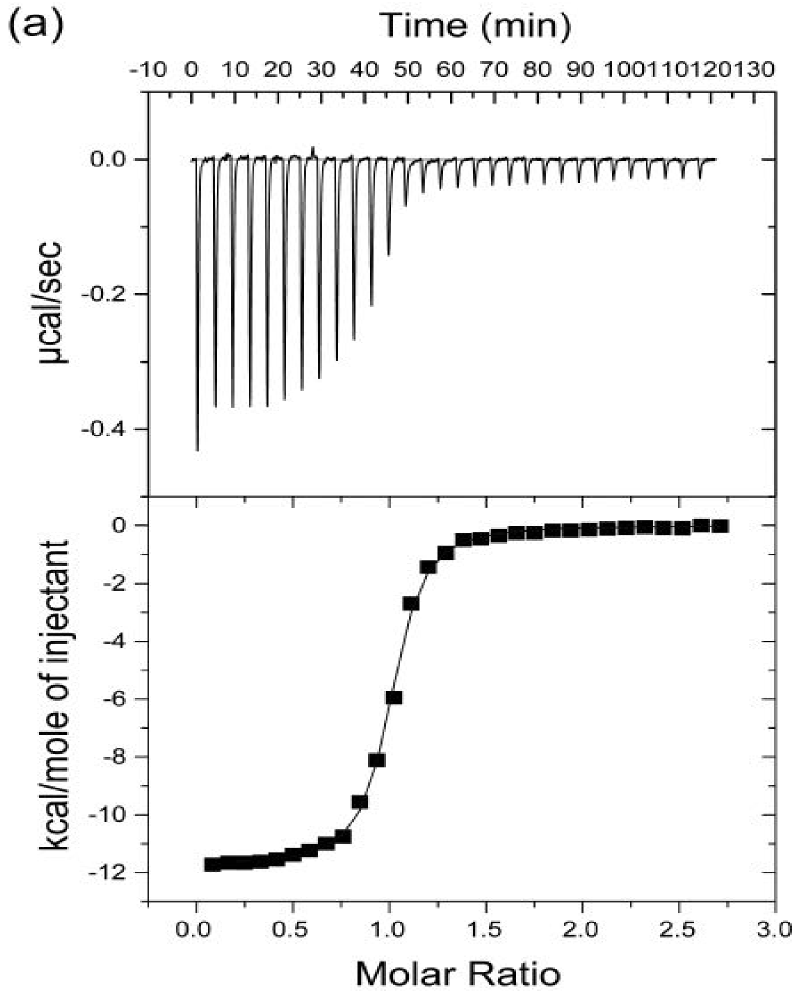


B. Differential scanning calorimetry（DSC）

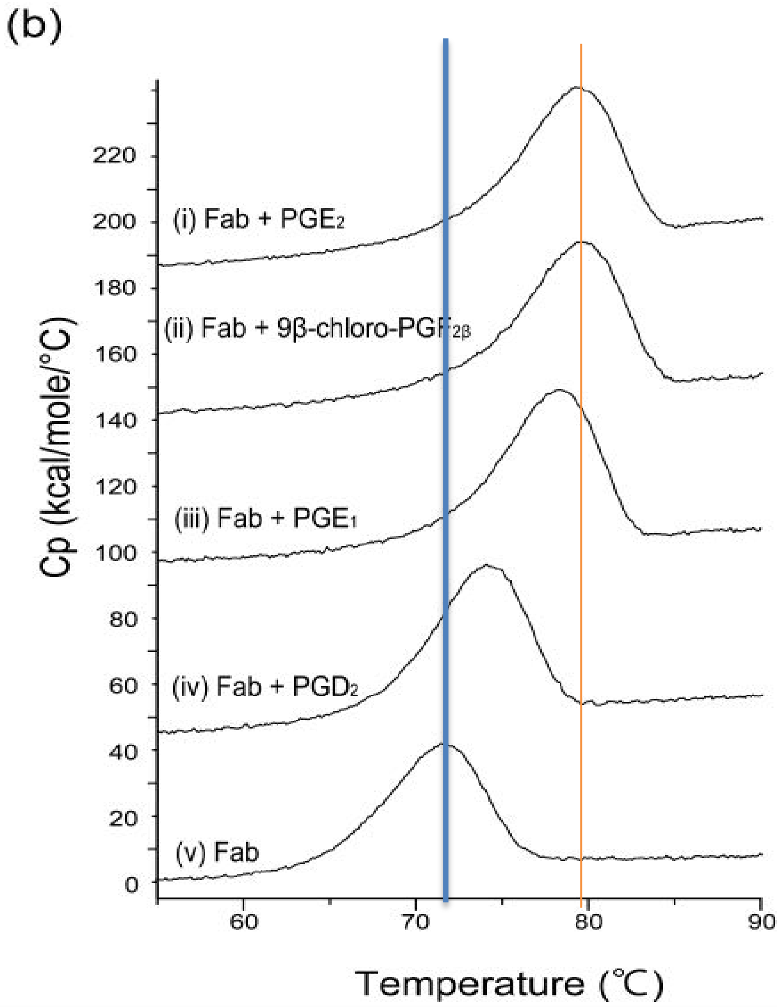

### Supplementary Fig. 2 A hydrogen bonding side chains on the $PGE_2$ binding crevasse interior.

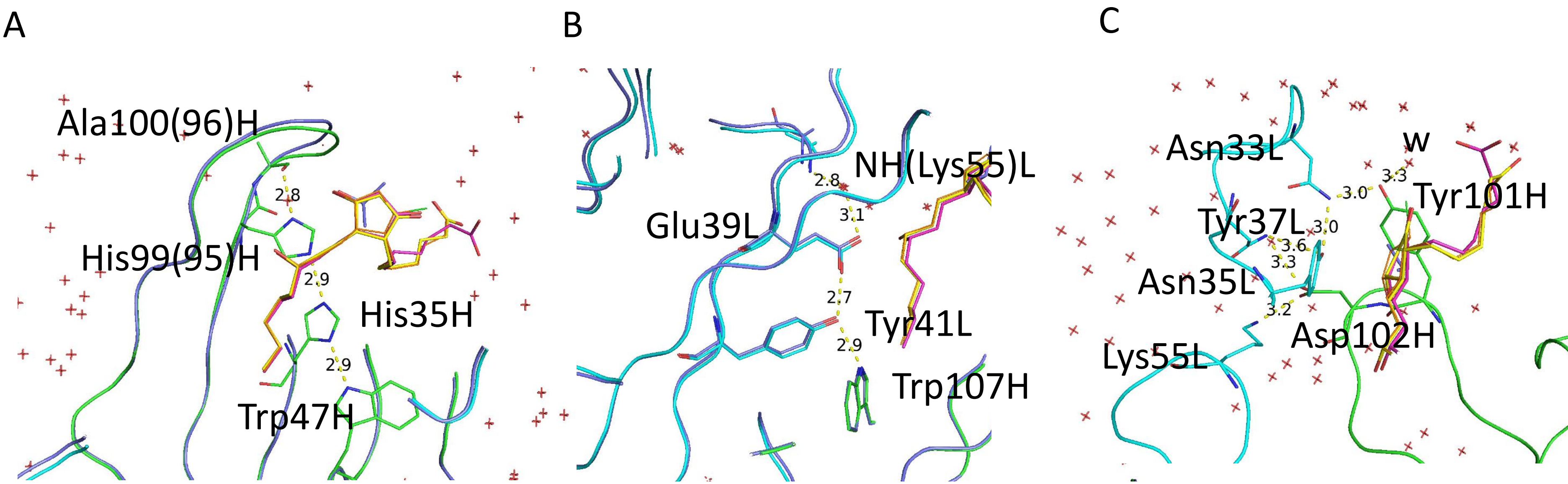

### Supplementary Figure 3 the $V_H$ domain shift between $PGE_2$ and $PGE_1$ complexed mAbPGE2 in stereo

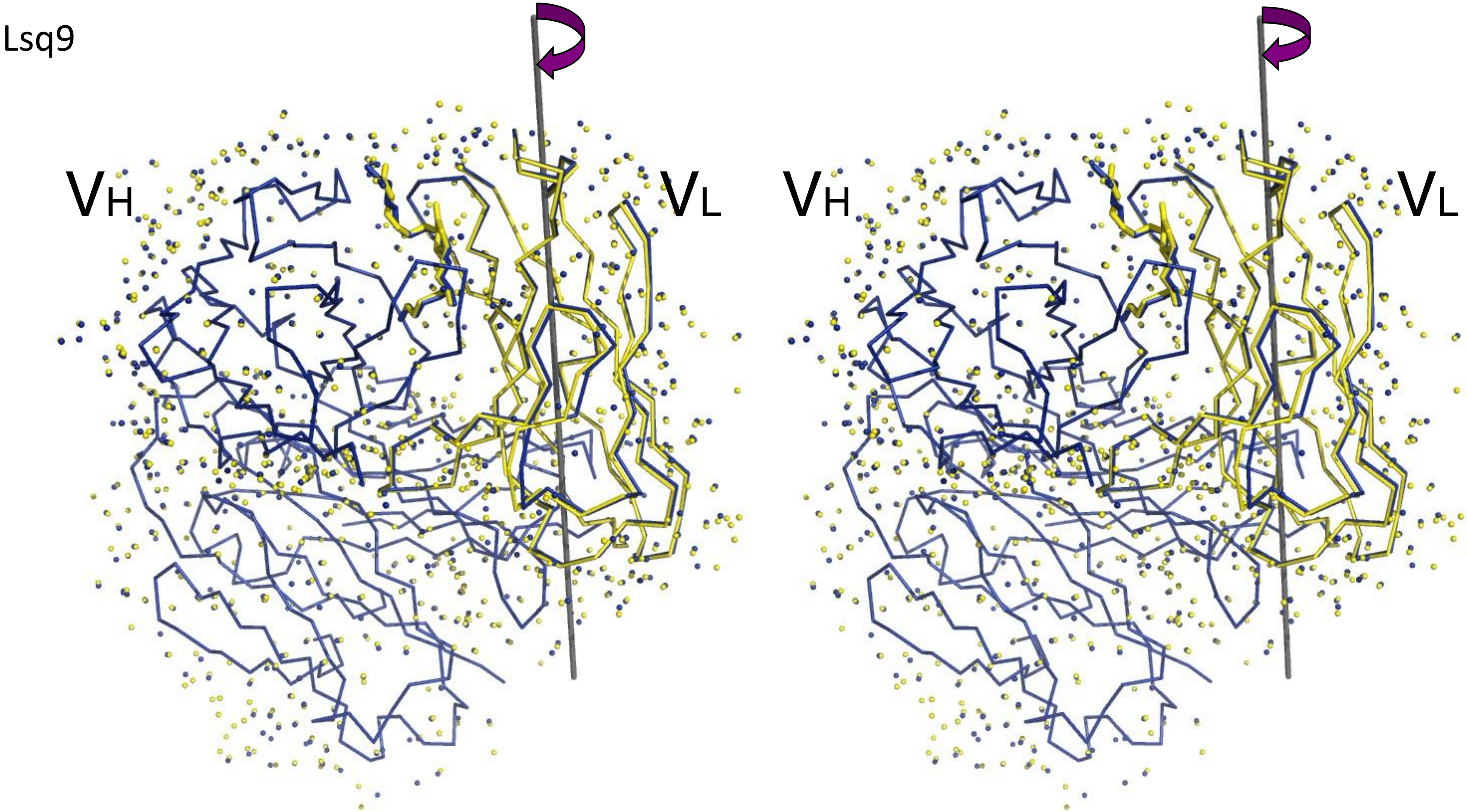

Supplementary Fig. 4 ITC measurements of binding reaction of mAb with $PGE_2$ and $PGE_1$ in BSA.

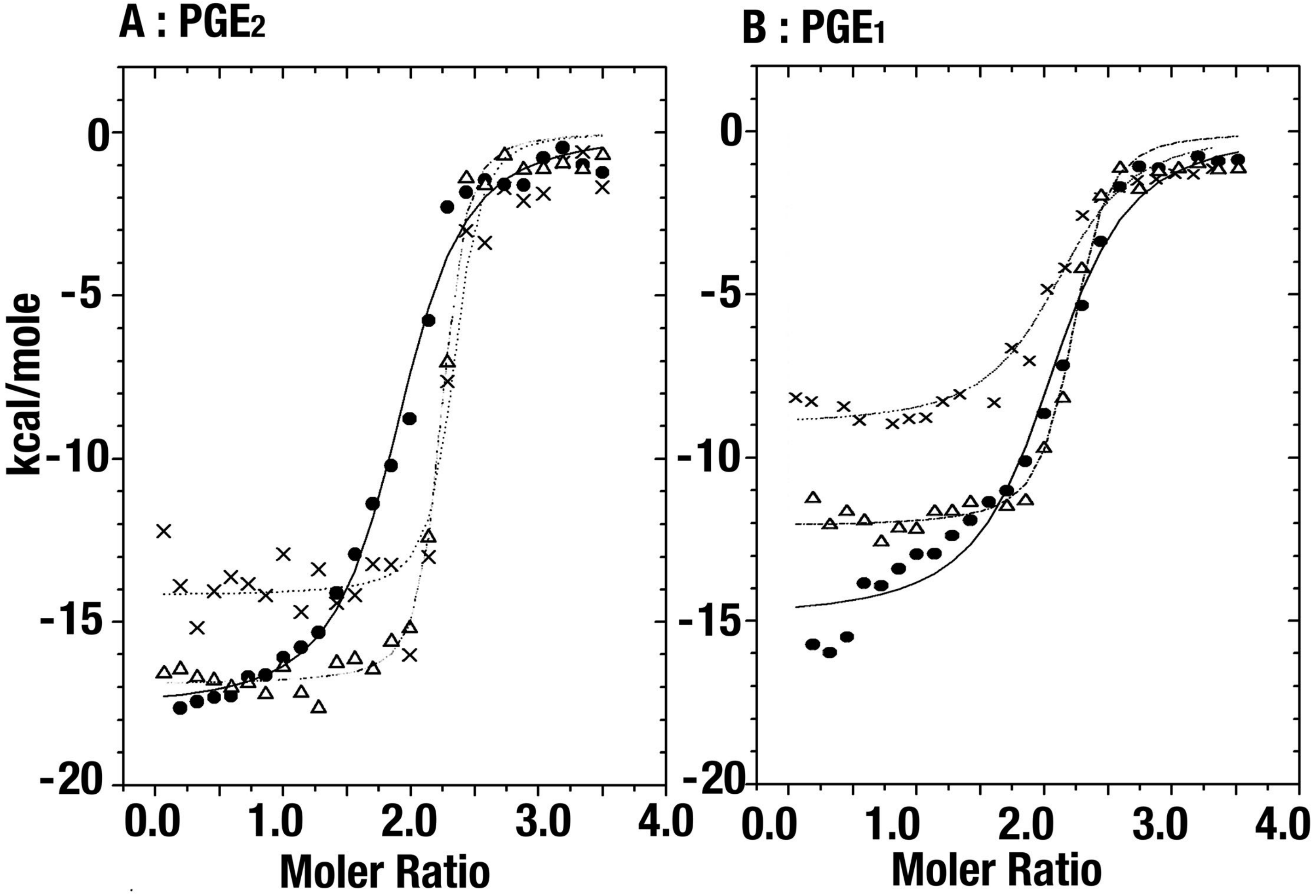


Supplementary Fig. 2 mAb ITC by antigen PG titrations of mAb, by $PGE_2$ (A) and $PGE_1$(B). -●- : 0 mg/mL BSA, --△--: 10 mg/mL BSA, ･･×･･: 100 mg/mL BSA

Supplementary Fig. 5 EIA of mAb with $PGE_2$ and PGs

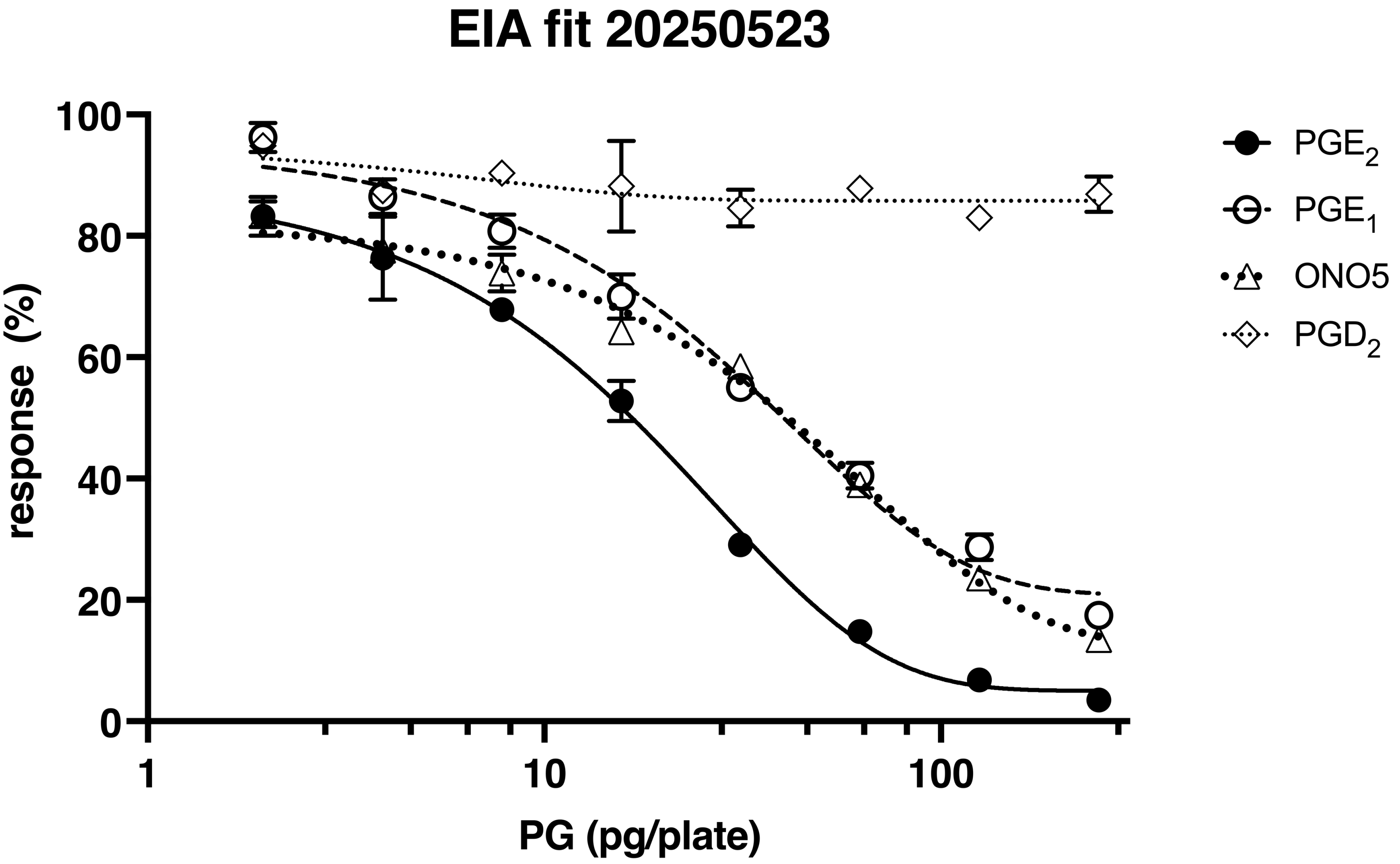

### Supplementary Fig. 6 Average B-factors of each VH, CH1, VL or CL of Fab:$PGE_2$, Fab:$PGE_1$ or Fab:ONO5 as well as antigen free.

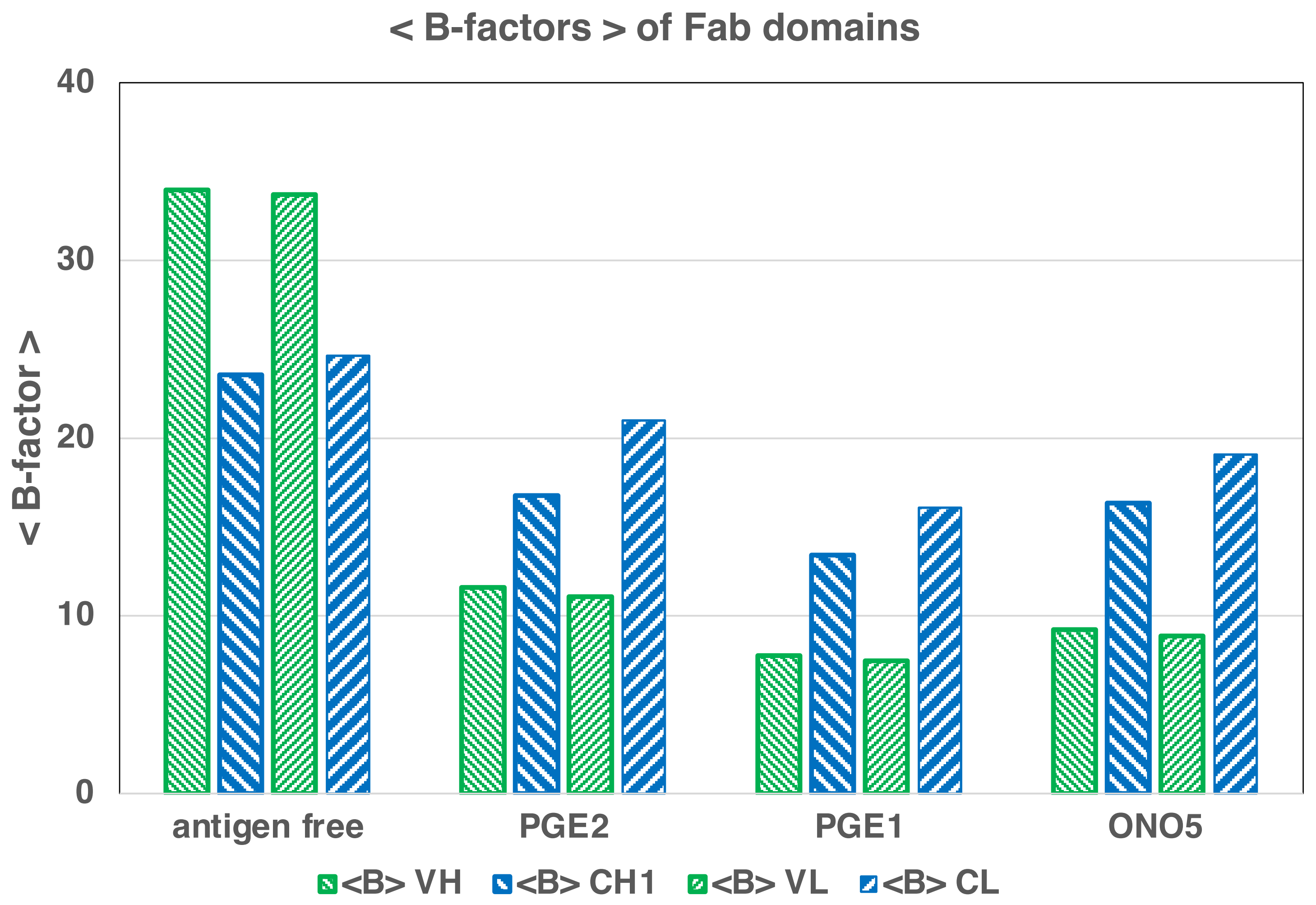

### Supplementary Fig. 7 Secondary Structure assignment of VH and VL sequences of Fab with Kabat and Chothia Numbering

#### Secondary Structure of antiPGE2 Fab in Kabat and Chothia Numberings

2025/11/12

**Light**

| | | | | | A0o | | | | | | Ai | | | | | | | | | | Bo | | | | | | | | | | | | | | | | | | | Ci |
|---|---|---|---|---|---|---|---|---|---|---|---|---|---|---|---|---|---|---|---|---|---|---|---|---|---|---|---|---|---|---|---|---|---|---|---|---|---|---|---|---|
| antiPGE2 Fab | 1 | 2 | 3 | 4 | 5 | 6 | 7 | 8 | 9 | 10 | 11 | 12 | 13 | 14 | 15 | 16 | 17 | 18 | 19 | 20 | 21 | 22 | 23 | 24 | 25 | 26 | 27 | 28 | 29 | 30 | 31 | 32 | 33 | 34 | 35 | 36 | 37 | 38 | 39 | 40 |
| Chothia | 1 | 2 | 3 | 4 | 5 | 6 | 7 | 8 | 9 | 10 | 11 | 12 | 13 | 14 | 15 | 16 | 17 | 18 | 19 | 20 | 21 | 22 | 23 | 24 | 25 | 26 | 27 | 28 | 29 | 30 | 30A | 30B | 30C | 30D | 30E | 31 | 32 | 33 | 34 | 35 |
| | D | V | L | M | T | Q | T | P | L | S | L | P | V | S | L | G | D | Q | A | S | I | S | C | R | S | S | Q | S | I | V | H | S | N | G | N | T | Y | L | E | W |
| Kabat | 1 | 2 | 3 | 4 | 5 | 6 | 7 | 8 | 9 | 10 | 11 | 12 | 13 | 14 | 15 | 16 | 17 | 18 | 19 | 20 | 21 | 22 | 23 | 24 | 25 | 26 | 27 | 27A | 27B | 27C | 27D | 27E | 28 | 29 | 30 | 31 | 32 | 33 | 34 | 35 |
| | | | | | | | | | | | | | | | | | | | | | | | | | | | | | | | | CDR1L | | | | | | | | |

| | | | | | | | | | | | | C'i | | | | | | C"i | | | | | | | | | | | Do | | | | | | | | Eo | | | |
|---|---|---|---|---|---|---|---|---|---|---|---|---|---|---|---|---|---|---|---|---|---|---|---|---|---|---|---|---|---|---|---|---|---|---|---|---|---|---|---|---|
| antiPGE2 Fab | 41 | 42 | 43 | 44 | 45 | 46 | 47 | 48 | 49 | 50 | 51 | 52 | 53 | 54 | 55 | 56 | 57 | 58 | 59 | 60 | 61 | 62 | 63 | 64 | 65 | 66 | 67 | 68 | 69 | 70 | 71 | 72 | 73 | 74 | 75 | 76 | 77 | 78 | 79 | 80 |
| Chothia | 36 | 37 | 38 | 39 | 40 | 41 | 42 | 43 | 44 | 45 | 46 | 47 | 48 | 49 | 50 | 51 | 52 | 53 | 54 | 55 | 56 | 57 | 58 | 59 | 60 | 61 | 62 | 63 | 64 | 65 | 66 | 67 | 68 | 69 | 70 | 71 | 72 | 73 | 74 | 75 |
| | Y | L | Q | K | P | G | Q | S | P | K | L | L | I | Y | K | V | S | N | R | F | S | G | V | P | D | R | F | S | G | S | G | S | G | T | D | F | T | L | K | I |
| Kabat | 36 | 37 | 38 | 39 | 40 | 41 | 42 | 43 | 44 | 45 | 46 | 47 | 48 | 49 | 50 | 51 | 52 | 53 | 54 | 55 | 56 | 57 | 58 | 59 | 60 | 61 | 62 | 63 | 64 | 65 | 66 | 67 | 68 | 69 | 70 | 71 | 72 | 73 | 74 | 75 |
| | | | | | | | | | | | | | | | | | CDR2L | | | | | | | | | | | | | | | | | | | | | | | |

| | | | | | | | | | | | | Fi | | | | | | | | | | | | | | | | | Gi | | | | | | | |
|---|---|---|---|---|---|---|---|---|---|---|---|---|---|---|---|---|---|---|---|---|---|---|---|---|---|---|---|---|---|---|---|---|---|---|---|---|
| antiPGE2 Fab | 81 | 82 | 83 | 84 | 85 | 86 | 87 | 88 | 89 | 90 | 91 | 92 | 93 | 94 | 95 | 96 | 97 | 98 | 99 | 100 | 101 | 102 | 103 | 104 | 105 | 106 | 107 | 108 | 109 | 110 | 111 | 112 | 113 | 114 | 115 | 116 |
| Chothia | 76 | 77 | 78 | 79 | 80 | 81 | 82 | 83 | 84 | 85 | 86 | 87 | 88 | 89 | 90 | 91 | 92 | 93 | 94 | 95 | 96 | 97 | 98 | 99 | 100 | 101 | 102 | 103 | 104 | 105 | 106 | 107 | 108 | 109 | 110 | 111 |
| | N | R | V | E | A | E | D | L | G | I | Y | Y | C | L | Q | G | S | H | V | P | L | T | F | G | A | G | T | T | L | E | L | K | R | A | - | - |
| Kabat | 76 | 77 | 78 | 79 | 80 | 81 | 82 | 83 | 84 | 85 | 86 | 87 | 88 | 89 | 90 | 91 | 92 | 93 | 94 | 95 | 96 | 97 | 98 | 99 | 100 | 101 | 102 | 103 | 104 | 105 | 106 | 107 | 108 | 109 | 110 | 111 |
| | | | | | | | | | | | | | | | | CDR3L | | | | | | | | | | | | | | | | | | | | |

- Outer Sheet
- Inner Sheet
- Helix

**Heavy**

| | | | | | A0o | | | | | | Ai | | | | | | | | | | Bo | | | | | | | | | Pivot point | | | | | | Ci | | | | |
|---|---|---|---|---|---|---|---|---|---|---|---|---|---|---|---|---|---|---|---|---|---|---|---|---|---|---|---|---|---|---|---|---|---|---|---|---|---|---|---|---|
| antiPGE2 Fab | 1 | 2 | 3 | 4 | 5 | 6 | 7 | 8 | 9 | 10 | 11 | 12 | 13 | 14 | 15 | 16 | 17 | 18 | 19 | 20 | 21 | 22 | 23 | 24 | 25 | 26 | 27 | 28 | 29 | 30 | 31 | 32 | 33 | 34 | 35 | 36 | 37 | 38 | 39 | 40 |
| Chothia | 1 | 2 | 3 | 4 | 5 | 6 | 7 | 8 | 9 | 10 | 11 | 12 | 13 | 14 | 15 | 16 | 17 | 18 | 19 | 20 | 21 | 22 | 23 | 24 | 25 | 26 | 27 | 28 | 29 | 30 | 31 | 32 | 33 | 34 | 35 | 36 | 37 | 38 | 39 | 40 |
| | K | V | Q | L | Q | Q | S | G | A | E | L | V | K | P | G | A | S | V | K | L | S | C | K | A | S | G | Y | T | F | T | E | Y | F | I | H | W | V | K | Q | R |
| Kabat | 1 | 2 | 3 | 4 | 5 | 6 | 7 | 8 | 9 | 10 | 11 | 12 | 13 | 14 | 15 | 16 | 17 | 18 | 19 | 20 | 21 | 22 | 23 | 24 | 25 | 26 | 27 | 28 | 29 | 30 | 31 | 32 | 33 | 34 | 35 | 36 | 37 | 38 | 39 | 40 |
| | | | | | | | | | | | | | | | | | | | | | | | | | | | | | | CDR1H | | | | | | | | | | |

| | | | | | | | | | C' | | | | | | | | | | C" | | | | | | | | | | | Do | | | | | | | | | | Eo |
|---|---|---|---|---|---|---|---|---|---|---|---|---|---|---|---|---|---|---|---|---|---|---|---|---|---|---|---|---|---|---|---|---|---|---|---|---|---|---|---|---|
| antiPGE2 Fab | 41 | 42 | 43 | 44 | 45 | 46 | 47 | 48 | 49 | 50 | 51 | 52 | 53 | 54 | 55 | 56 | 57 | 58 | 59 | 60 | 61 | 62 | 63 | 64 | 65 | 66 | 67 | 68 | 69 | 70 | 71 | 72 | 73 | 74 | 75 | 76 | 77 | 78 | 79 | 80 |
| Chothia | 41 | 42 | 43 | 44 | 45 | 46 | 47 | 48 | 49 | 50 | 51 | 52 | 52A | 53 | 54 | 55 | 56 | 57 | 58 | 59 | 60 | 61 | 62 | 63 | 64 | 65 | 66 | 67 | 68 | 69 | 70 | 71 | 72 | 73 | 74 | 75 | 76 | 77 | 78 | 79 |
| | S | G | Q | G | L | E | W | I | G | W | F | Y | P | G | S | G | S | L | N | Y | N | G | K | F | K | D | K | A | T | F | T | A | D | K | S | S | S | T | V | Y |
| Kabat | 41 | 42 | 43 | 44 | 45 | 46 | 47 | 48 | 49 | 50 | 51 | 52 | 52A | 53 | 54 | 55 | 56 | 57 | 58 | 59 | 60 | 61 | 62 | 63 | 64 | 65 | 66 | 67 | 68 | 69 | 70 | 71 | 72 | 73 | 74 | 75 | 76 | 77 | 78 | 79 |
| | | | | | | | | | | | | | | | | CDR2H | | | | | | | | | | | | | | | | | | | | | | E | | |

| | | | | | | | | | | | | | | | Fi | | | | | | | | | | | | | | | | | | | | | | | |
|---|---|---|---|---|---|---|---|---|---|---|---|---|---|---|---|---|---|---|---|---|---|---|---|---|---|---|---|---|---|---|---|---|---|---|---|---|---|---|
| antiPGE2 Fab | 81 | 82 | 83 | 84 | 85 | 86 | 87 | 88 | 89 | 90 | 91 | 92 | 93 | 94 | 95 | 96 | 97 | 98 | 99 | 100 | 101 | 102 | 103 | 104 | 105 | 106 | 107 | 108 | 109 | 110 | 111 | 112 | 113 | 114 | 115 | 116 | 117 | 118 |
| Chothia | 80 | 81 | 82 | 82A | 82B | 82C | 83 | 84 | 85 | 86 | 87 | 88 | 89 | 90 | 91 | 92 | 93 | 94 | 95 | 96 | 97 | 98 | 99 | 100 | 101 | 102 | 103 | 104 | 105 | 106 | 107 | 108 | 109 | 110 | 111 | 112 | 113 | 114 |
| | L | E | L | S | R | L | T | S | E | D | S | A | V | Y | F | C | A | S | H | A | Y | D | K | E | P | Y | W | G | Q | G | T | L | V | T | V | S | A | - |
| Kabat | 80 | 81 | 82 | 82A | 82B | 82C | 83 | 84 | 85 | 86 | 87 | 88 | 89 | 90 | 91 | 92 | 93 | 94 | 95 | 96 | 97 | 98 | 99 | 100 | 101 | 102 | 103 | 104 | 105 | 106 | 107 | 108 | 109 | 110 | 111 | 112 | 113 | 114 |
| | | | | | | | | | | | | | | | | | | | | | | CDR3H | | | | | | | | | | | | | | | | |

### Supporting Appendix 1: Preparation and characterizations of mAb and Fab

#### 12.5 % SDS-Page of mAb and Fab

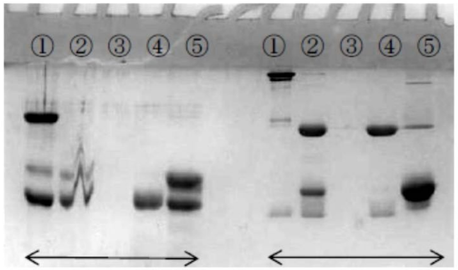


**β-mel. ( + )** **( - )**

① intact mAb
② papain digestion filter through
③ Papain digestion concentrate
④ Mono-Q Void
⑤ Mono-Q Fraction

#### (1) Cation chromatography

monoS 5/50 (1ml)
Frow rate :1 ml/min
Buffer: A:20mM Na phosphate 5.5, B:A+ 1M NaCl
Elution: 0↑20%B (20 CV gradient )

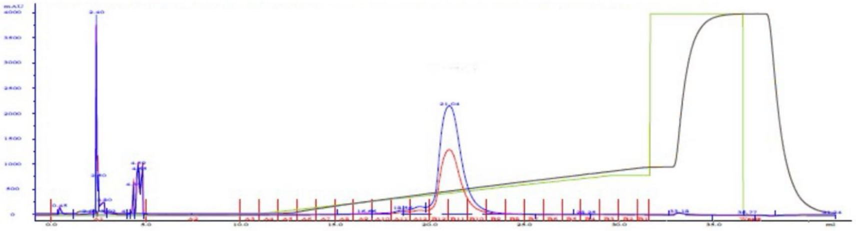

#### (2) Anion chromatography

monoQ 5/50 (1ml)
Frow rate :1 ml/min
Buffer: A:25mM Tris 8.0, 25mM NaCl B: 25mM Tris 8.0, 1M NaCl
Elution: 0↑100%B (0CV)

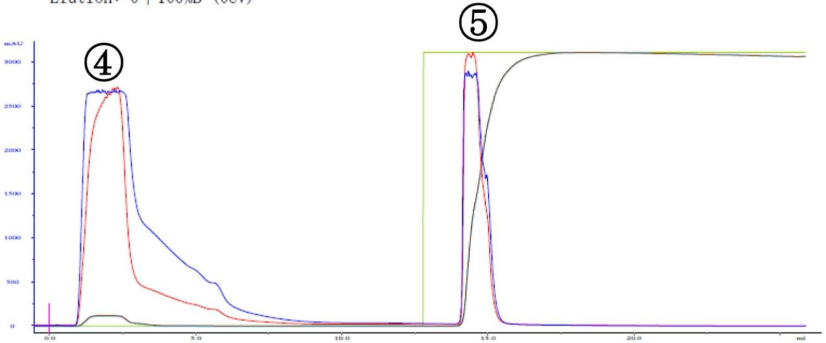


#### (3) Gel filtration chromatography

superdex 200 (24ml)
Frow rate : 0.5 ml/min
Buffer: 10mM Na phosphate pH7.0, 140mM NaCl, 5% 2-propanol
Sample collected (Arrow)

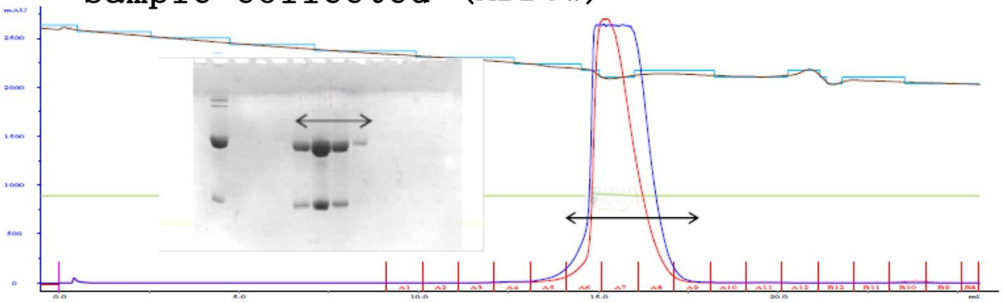

### Supporting Appendix 1’: Conventional Large Preparation of anti-$PGE_2$ Fab (13)

【Production IgG with no serum culture～ gel filtration chromatography 】
Hybridoma cell culture media
↓
【Protein A chromatography】
↓0.1M Na-Citrate pH4.0 elution buffer
↓neutriazed by 10% 1M Tris pH9.0
↓ yield: ca. 7-8mg per 400ml clture
【dialysis】PBS 4℃ 2 hrs
↓
【contration】> 5mg/ml
↓
【desalt】
↓75mM NaCl, 2mM EDTA, 5mM azide
↓75mM Na-Phosphate pH7.0
【mAb digention】37℃ 3 hrs
↓mAb 5mg/ml 1ml
↓papain 0.5mg/ml 10μl
↓cysteine 1M 20μl
Termination
with 100μl 0.2M *N*-methymalaimide
↓
【ProteinG chromatography】adsorb
↓0.1M Glycine HCl pH2.75 elution buffer
↓ neutriazed by 10% 1M Tris pH9.0
【concentration and desalting column】
↓PBS pH7.0
↓ + 2 eq. molar $PGE_2$
【Gel filtration hromatography】
↓PBS pH7.0
【concentration 】
↓ > 20mg/ml
↓ Yield ca. 3mgbFab per 10mg mAb
【crystallization】

### Supporting Appendix 2: SDS PAGE and partial amino acid sequences of the purified Papain digested anti-$PGE_2$ mouse IgGκ with protein A and G.

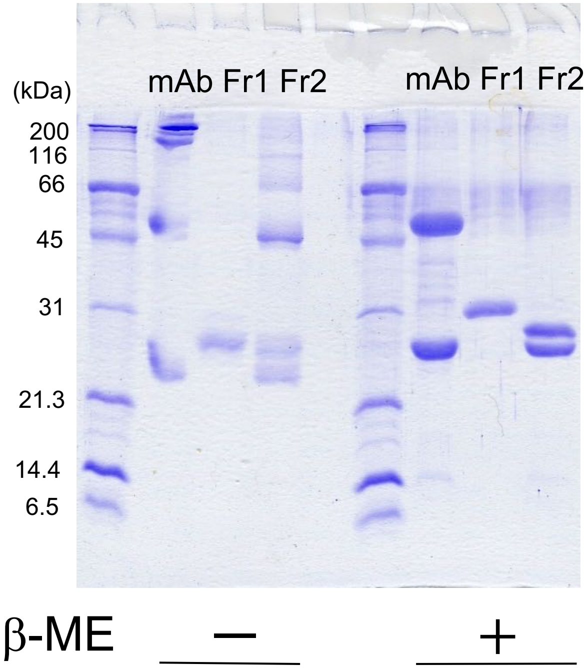


**N-terminal sequences of Edman degradation**

```
Fr.1:TVPEVSSVFI  (Heavy Chain 260-)
Fr.2:KVQLQQSGAE  (Heavy Chain 33-)
    +DVLMTQTPLS  (Light Chain 37-)
```

**LC-MS/MS fragments of n gel trypsin digestions pepttides of anti-PGE2 mouse IgG**

Fr. 1 :Upper panel and and Fr. 2:lowere panel.

| Chain | Residues | Sequence | Expected mass [M+H] $^{+1}$ *m/z* | Observed mass *m/z* | Observed charge state |
|---|---|---|---|---|---|
| heavy | [260–275] | TVPEVSSVFIFPPKPK | 1771.999 | 886.72 | +2 |
| heavy | [276–285] | DVLTITLTPK | 1100.656 | 550.73 | +2 |
| heavy | [329–344] | SVSELPIMHQDWLNGK | 1853.921 | 927.94 | +2 |
| heavy | [372–382] | APQVYTIPPPK | 1210.683 | 605.87 | +2 |
| heavy | [420–436] | NTQPIMDTDGSYFVYSK | 1965.89 | 983.91 | +2 |
| heavy | [437–441] | LNVQK | 601.367 | 301.15 | +2 |

| Chain | Residues | Sequence | Expected mass [M+H] $^{+1}$ *m/z* | Observed mass *m/z* | Observed charge state |
|---|---|---|---|---|---|
| heavy | [ 57– 71] | ASGYTFTEYFIHWVK | 1848.896 | 925.48 | +2 |
| heavy | [ 74– 96] | SGQGLEWIGWFYPGSGSLNYNGK | 2517.183 | 840.16 | +3 |
| heavy | [108–118] | SSSTVYLELSR | 1241.637 | 621.52 | +2 |
| heavy | [137–152] | EPYWGQGTLVTVSAAK | 1706.875 | 853.90 | +2 |
| Light | [103–115] | FSGSGSGTDFTLK | 1303.616 | 652.54 | +2 |
| Light | [197–210] | QNGVLNSWTDQDSK | 1591.735 | 796.95 | +2 |
| Light | [211–224] | DSTYSMSSTLTLTK | 1534.731 | 768.15 | +2 |

### Supporting Appendix 3: Anti-$PGE_2$ immunoglobin cDNA and deduced amino acid with partial amino acid sequence of Heavy chain of papain digested fragments as Fab and Fc. (Fraction 1 and Fraction2).

```
                                   acgcggggacccatgatcagtatcctctccacagtcactgaacacactgactcaaacc

Fr.2-1   atg gaa tgg tgc tgg gtc ttt ctc ttc ctc ctg tca gta act gca ggt gtc cac tcc AAG    F_H
Fab_H -19 M   E   W   C   W   V   F   L   F   L   L   S   V   T   A   G   V   H   S   K      1
         GTC CAG CTG CAG CAG TCT GGA GCT GAG TTG GTG AAA CCC GGG GCA TCA GTG AAG CTG TCC
     2    V   Q   L   Q   Q   S   G   A   E   L   V   K   P   G   A   S   V   K   L   S     21
         TGC AAG GCT TCT GGC TAC ACC TTC ACT GAG TAT TTT ATA CAC TGG GTA AAG CAG AGG TCT
    22    C   K   A   S   G   Y   T   F   T   E   Y   F   I   H   W   V   K   Q   R   S     41
         GGA CAG GGT CTT GAG TGG ATT GGG TGG TTT TAC CCT GGA AGT GGT AGT TTA AAT TAC AAT
    42    G   Q   G   L   E   W   I   G   W   F   Y   P   G   S   G   S   L   N   Y   N     61
         GGG AAA TTC AAG GAC AAG GCC ACA TTT ACT GCG GAC AAA TCC TCC AGC ACA GTC TAT TTG
    62    G   K   F   K   D   K   A   T   F   T   A   D   K   S   S   S   T   V   Y   L     81
         GAG CTT AGT AGA TTG ACA TCT GAA GAC TCT GCG GTC TAT TTC TGT GCA AGC CAC GCT TAC
    82    E   L   S   R   L   T   S   E   D   S   A   V   Y   F   C   A   S   H   A   Y    101
         GAC AAG GAG CCG TAC TGG GGC CAA GGG ACT CTG GTC ACT GTC TCT GCA GCC AAA ACG ACA
   102    D   K   E   P   Y   W   G   Q   G   T   L   V   T   V   S   A   A   K   T   T    121
         CCC CCA TCT GTC TAT CCA CTG GCC CCT GGA TCT GCT GCC CAA ACT AAC TCC ATG GTG ACC
   122    P   P   S   V   Y   P   L   A   P   G   S   A   A   Q   T   N   S   M   V   T    141
         CTG GGA TGC CTG GTC AAG GGC TAT TTC CCT GAG CCA GTG ACA GTG ACC TGG AAC TCT GGA
   142    L   G   C   L   V   K   G   Y   F   P   E   P   V   T   V   T   W   N   S   G    161
         TCC CTG TCC AGC GGT GTG CAC ACC TTC CCA GCT GTC CTG CAG TCT GAC CTC TAC ACT CTG
   162    S   L   S   S   G   V   H   T   F   P   A   V   L   Q   S   D   L   Y   T   L    181
         AGC AGC TCA GTG ACT GTC CCC TCC AGC ACC TGG CCC AGC GAG ACC GTC ACC TGC AAC GTT
   182    S   S   S   V   T   V   P   S   S   T   W   P   S   E   T   V   T   C   N   V    201
         GCC CAC CCG GCC AGC AGC ACC AAG GTG GAC AAG AAA ATT GTG ccc agg gat tgt ggt tgt
   202    A   H   P   A   S   S   T   K   V   D   K   K   I   V   P   R   D   C   G   C    218+3
Fr.1     aag cct tgc ata tgt aca gtc cca gaa gta tca tct gtc ttc atc ttc ccc cca aag ccc
Fc        K   P   C   I   C   T   V   P   E   V   S   S   V   F   I   F   P   P   K   P    241
         AAG GAT GTG CTC ACC ATT ACT CTG ACT CCT AAG GTC ACG TGT GTT GTG GTA GAC ATC AGC
   242    K   D   V   L   T   I   T   L   T   P   K   V   T   C   V   V   V   D   I   S    261
         AAG GAT GAT CCC GAG GTC CAG TTC AGC TGG TTT GTA GAT GAT GTG GAG GTG CAC ACA GCT
   262    K   D   D   P   E   V   Q   F   S   W   F   V   D   D   V   E   V   H   T   A    281
         CAG ACG CAA CCC CGG GAG GAG CAG TTC AAC AGC ACT TTC CGC TCA GTC AGT GAA CTT CCC
   282    Q   T   Q   P   R   E   E   Q   F   N   S   T   F   R   S   V   S   E   L   P    301
         ATC ATG CAC CAG GAC TGG CTC AAT GGC AAG GAG TTC AAA TGC AGG GTC AAC AGT GCA GCT
   302    I   M   H   Q   D   W   L   N   G   K   E   F   K   C   R   V   N   S   A   A    321
         TTC CCT GCC CCC ATC GAG AAA ACC ATC TCC AAA ACC AAA GGC AGA CCG AAG GCT CCA CAG
   322    F   P   A   P   I   E   K   T   I   S   K   T   K   G   R   P   K   A   P   Q    341
         GTG TAC ACC ATT CCA CCT CCC AAG GAG CAG ATG GCC AAG GAT AAA GTC AGT CTG ACC TGC
   342    V   Y   T   I   P   P   P   K   E   Q   M   A   K   D   K   V   S   L   T   C    361
         ATG ATA ACA GAC TTC TTC CCT GAA GAC ATT ACT GTG GAG TGG CAG TGG AAT GGG CAG CCA
   362    M   I   T   D   F   F   P   E   D   I   T   V   E   W   Q   W   N   G   Q   P    381
         GCG GAG AAC TAC AAG AAC ACT CAG CCC ATC ATG GAC ACA GAT GGC TCT TAC TTC GTC TAC
   382    A   E   N   Y   K   N   T   Q   P   I   M   D   T   D   G   S   Y   F   V   Y    401
         AGC AAG CTC AAT GTG CAG AAG AGC AAC TGG GAG GCA GGA AAT ACT TTC ACC TGC TCT GTG
   402    S   K   L   N   V   Q   K   S   N   W   E   A   G   N   T   F   T   C   S   V    421
         TTA CAT GAG GGC CTG CAC AAC CAC CAT ACT GAG AAG AGC CTC TCC CAC TCT CCT GGT AAA
   422    L   H   E   G   L   H   N   H   H   T   E   K   S   L   S   H   S   P   G   K    441
         tccctgtataaataaagcacccagcactgccttgggaaaaaaaaaaaaaaaaaaaaaaaaaaaaagtactctgcgttga

         taccactgcttgccctatagtgagtcgtattag
```

**Supplementary Fig. XX: Amino acid sequencing of the papain digested $PGE_2$ mAb in protein-G chromatography**

*N*-terminal sequencing sequences with a automated Edman-sequencer (Red letter with Red arrow) and Liquid chromatography MS/MS sequencing of the trypsin-digested peptides (Skyblue letter with Skyblue line) of the Fractions 1 and 2 in the protein-G chromatography of papain-proteolyzed PGE2 mAb.

> Fab $V_H$-$F_C$1 Heavy chain in Fr.2

KVQLQQSGAELVKPGASVKLSCKASGYTFTEYFIHWVKQRSGQGLEWIGWFYPGSGSLNYNGKFKDKATFTADKSSST

VYLELSRLTSEDSAVYFCASHAYDKEPYWGQGTLVTVSAAKTTPPSVYPLAPGSAAQTNSMVTLGCLVKGYFPEPVTV

TWNSGSLSSGVHTFPAVLQSDLYTLSSSVTVPSSTWPSETVTCNVAHPASSTKVDKKIVPRDC*GCKPCIC*

>Fab $V_L$-$F_L$ Light chain in Fr.2

mklpvrllvlmfwipasssDVLMTQTPLSLPVSLGDQASISCRSSQSIVHSNGNTYLEWYLQKPGQSPKLLIYKVSNR

FSGVPDRFSGSGSGTDFTLKINRVEAEDLGIYYCLQGSHVPLTFGAGTTLELKRADAAPTVSIFPPSSEQLTSGGASV

VCFLNNFYPKDINVKWKIDGSERQNGVLNSWTDQDSKDSTYSMSSTLTLTKDEYERHNSYTCEATHKTSTSPIVKSFN

RN*EC*

> Fc $F_{CH}$-2/$F_{CH}$-3 Heavy Chain in Fr. 1

TVPEVSSVFIFPPKPKDVLTITLTPKVTCVVVDISKDDPEVQFSWFVDDVEVHTAQTQPREEQFNSTFRSVSELPIMH

QDWLNGKEFKCRVNSAAFPAPIEKTISKTKGRPKAPQVYTIPPPKEQMAKDKVSLTCMITDFFPEDITVEWQWNGQPA

ENYKNTQPIMDTDGSYFVYSKLNVQKSNWEAGNTFTCSVLHEGLHNHHTEKSLSHSPGK

### Supporting Appendix 4: Sequence alignment of VL and VH in anti-$PGE_2$ mAb and related mAbs.

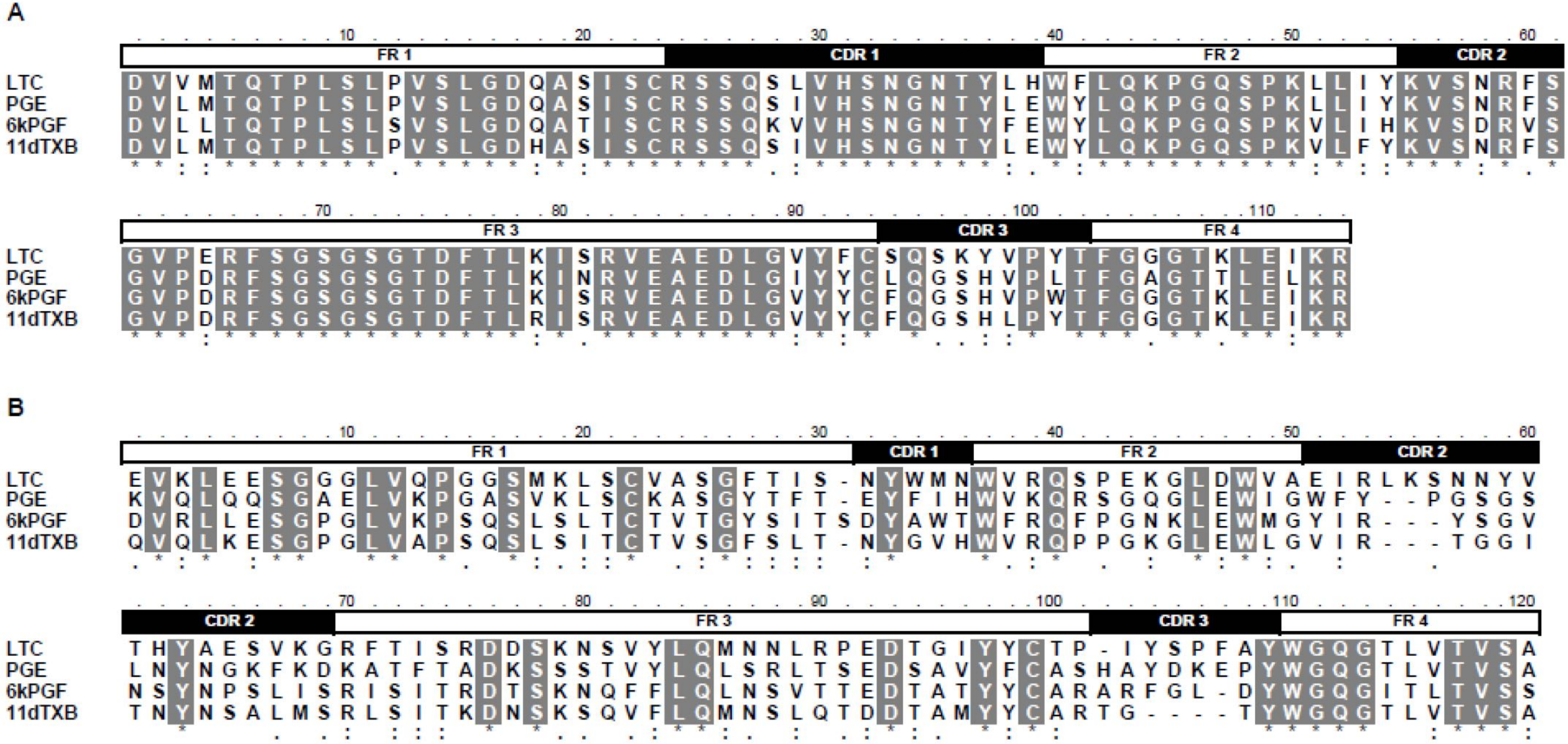


A: VL. B: VH. CDR are based on Kabat numbering.

Wu TT, Kabat EA. An analysis of the sequences of the variable regions of Bence Jones proteins and myeloma light chains and their implications for antibody complementarity. J Exp Med. 1970 Aug 1;132(2):211-50. doi: 10.1084/jem.132.2.211. PMID: 5508247; PMCID: PMC2138737.
